# Using synthetic MR images for distortion correction

**DOI:** 10.1101/2021.03.13.435270

**Authors:** David F. Montez, Andrew N. Van, Ryland L. Miller, Nicole A. Seider, Scott Marek, Annie Zheng, Dillan J. Newbold, Kristen Scheidter, Eric Feczko, Anders J. Perrone, Oscar Miranda-Dominguez, Eric A. Earl, Benjamin P. Kay, Abhinav K. Jha, Aristeidis Sotiras, Timothy O. Laumann, Deanna J. Greene, Evan M. Gordon, M. Dylan Tisdall, Andre van der Kouwe, Damien A. Fair, Nico U.F. Dosenbach

## Abstract

Functional MRI (fMRI) data acquired using echo-planar imaging (EPI) are highly distorted by magnetic field inhomogeneities. Distortion combined with underlying differences in image contrast between EPI and T1-weighted and T2-weighted (T1w/T2w) structural images makes the alignment of functional and anatomical images a challenge. Typically, separately acquired field map data are used to correct fMRI distortions and a flexible cost function insensitive to cross-modal differences in image contrast and intensity is used for aligning fMRI and anatomical images. The quality of alignment achieved with this approach can vary greatly and depends on the quality of field map data. In addition, many publicly available datasets lack field map data entirely. To address this issue, we developed *Synth*, a software package for distortion correction and cross-modal image registration that does not require separately acquired field map data. *Synth* combines information from T1w and T2w anatomical images to construct an idealized undistorted synthetic image that has similar contrast properties to fMRI data. The undistorted synthetic image then serves as an effective reference for individual-specific nonlinear unwarping to correct fMRI distortions. We demonstrate, in both pediatric (ABCD: Adolescent Brain Cognitive Development) and adult (MSC: Midnight Scan Club) data that *Synth* performs comparably well to other leading distortion correction approaches that utilize field map data, and often outperforms them. Field map-less distortion correction with *Synth* allows accurate and precise registration of fMRI data with missing or corrupted field map information.

## 1. Introduction

BOLD-weighted (blood-oxygenation level dependent) functional MRI (fMRI) data obtained using echo planar imaging (EPI) is severely distorted by inhomogeneities affecting the primary magnetic field [1, 2, 3, 4, 5]. EPI distortion, which consists of localized spatial deformation and loss of BOLD signal intensity, prominently affects areas containing large local differences in magnetic susceptibility. Due to the apposition of diamagnetic tissue and paramagnetic air in the sinuses and ear canals, regions of the image containing the orbitofrontal cortex and the inferior temporal lobes often suffer the most severe distortions. The susceptibility-induced artifacts are spatially non-uniform and therefore interfere with the performance of registration algorithms used during fMRI preprocessing to bring BOLD EPI images into alignment with their associated anatomical images (T1w, T2w).

Establishing the correspondence between brain anatomy and function is an important component of interpreting neuroimaging findings. Regardless of study design or choice of analysis space, registration and removal of EPI distortion and registration to T1w/T2w anatomical images are crucial steps in the analysis of fMRI data. For example, poor alignment counteracts the benefits of analyses performed with reference to participant-specific anatomy. Improper alignment of BOLD and anatomical images also degrades the performance of procedures that project volumetric fMRI data onto mesh surfaces derived from tissue segmentation [6]. The effects of poor registration and distortion correction also carry forward into group analyses. In group studies, anatomical images from many participants are separately aligned to a reference atlas. However, suboptimal alignment between a participant’s BOLD and anatomical images will propagate as nuisance variability that negatively affects group-level statistics for both task and resting state analyses [7, 8].

Significant effort has been devoted to developing methods to correct distortions affecting fMRI data. Currently, two primary methods are commonly used, both of which involve acquiring field map data at scan time. The first method involves acquiring a pair of EPI images with opposing phase encoding directions. Because the largest EPI distortions occur in the phase encoding direction, reversing the phase encoding direction also reverses the direction of the EPI distortions. By combining information from both directions, a displacement field that corrects for the underlying EPI distortion can be constructed [2]. The second field map method relies on the linear relationship between the phase of gradient echo data, local magnetic field inhomogeneities, and echo time. By recording MR data with two different echo times, a displacement field can be constructed that corrects for distortion caused by inhomogeneity [4]. The end product of each of these approaches is a nonlinear warp that indicates how each voxel must be displaced in order to correct for the image distortion.

Without mitigation, the validity and quality of corrections produced by either of the standard field map methods hinges on several practical points: First, subject movement during the acquisition of the field map data or during the time between the acquisition of the field map data and the corresponding EPI data can negatively affect corrections in two ways: 1) motion can reduce the quality of the field map data itself by introducing artifacts which reduce the accuracy of corrections; and 2) movement may also change the spatial structure of the distortion so that the geometry of the brain images differs in the field map data and the EPI data [9]. This may reduce the accuracy of the corrections simply by reducing the accuracy of the alignments between EPI data and field map data meant to provide the correction. Second, research has shown that dual-echo and phase-reversal methods for estimating EPI distortion perform differently across brain regions and that the method used to generate the echoes (spin-echo vs. gradient-echo) used in field map acquisitions can affect the accuracy of distortion correction [10]. Finally, the process of estimating the distortion correction from field map data, in most cases, relies on internal regularization parameters of the algorithm used to estimate the distortion. These parameters affect the overall smoothness of the final correcting warp, and may not be optimal for a particular set of image acquisition parameters or level of image detail.

The need to acquire field maps introduces a crucial failure point during data acquisition. Failure to acquire valid field map data can disqualify an entire dataset from inclusion in an analysis. This problem is further amplified in study designs requiring multiple field maps be collected across multiple sessions or even within sessions. In these cases, each field map represents a potential failure point where poor data collection can affect the quality of post-acquisition analysis. For instance, within the ABCD Annual Release 2.0 dataset (DOI 10.15154/1503209) 341 participants were missing valid field map data. At the time of writing, of the top five most downloaded fMRI datasets available on OpenNeuro.org (Flanker task (event-related), UCLA Consortium for Neuropsychiatric Phenomics LA5c Study, Classification learning, Forrest Gump, Multisubject, multimodal face processing), only one includes field map data. An effective implementation of a field map-less approach is highly desirable because it would allow for the correction of images for which field map information was either missing, corrupted, or never collected.

Motivated by the limitations of field maps and the potential to reinvestigate datasets lacking field maps, fMRI researchers have explored direct mapping approaches in which EPI distortion is corrected by non-linearly registering a participant’s distorted BOLD images directly to their undistorted anatomical image. Early implementations of direct mapping demonstrated that the approach could reduce distortion and improve global measures of image similarity (e.g., mutual information or squared-error) between BOLD and anatomical images [11, 12, 13, 14]. However, further research revealed that global measures of image similarity can be an unreliable indicator of nonrigid image alignment quality [15]. More recent implementations of the direct mapping approach include external group average field map data as a constraint on possible solutions (e.g., fieldmap-less SyN-Susceptibility Distortion Correction (SDC)). Quality assessment of distortion corrections produced by direct mapping demonstrate that final image alignment quality can be unreliable and tends to perform more poorly than high quality corrections using a field map [16, 17, 18, 19]. One likely cause for the variable performance of this approach is the fact that T1w, T2w, and BOLD images look quite different from one another. Because the average signal intensities associated with the tissues and fluids comprising the brain vary considerably across acquisition parameters, it is difficult to construct registration cost functions that can accurately reflect the true error introduced by EPI distortion [20].

Image correction using undistorted synthetic image references is another more recent approach to field map-less distortion correction. This strategy entails the estimation of an undistorted auxiliary EPI image that serves as a reference for unwarping. The Synb0-DisCo algorithm is a notable example of this approach that employs trained deep learning neural networks to transform undistorted T1w images into a synthetic EPI image which is used as an input image for FSL’s *topup* algorithm [21]. Initial assessments of this approach suggest that it can potentially produce distortion corrections that are comparable to those produced by high quality field map data. However, because T1w and EPI image contrast can vary significantly across acquisitions and depends on the accuracy of the intensity bias correction applied to the images, it is unclear how well these trained deep learning approaches will perform on arbitrary datasets. In addition, there is potential for synthetic images produced by the deep learning networks to include spurious image artifacts which may affect the reliability of distortion corrections in practice [22, 23].

We sought to combine direct mapping and synthetic reference image approaches to produce a more reliable field map-less distortion correction algorithm -one that does not rely on priors established by training data. To accomplish this, we developed a modeling framework that allows us to combine information from a participant’s T1w, T2w, and BOLD images in order to construct a synthetic image that has the contrast properties of a BOLD image and the undistorted geometry and high resolution (∼ 1 mm^3^) of typical T1w/T2w images. We hypothesized that these synthetic BOLD images would serve as ideal targets for estimating field map corrections with currently available nonlinear warping software, and thereby improve direct mapping quality. Synthetic reference images created in this way do not rely on trained models. Consequently, this approach flexibly adapts to the particular contrast properties of a given dataset and may be less prone to the influence of deep learning network artifacts.

Here, we detail *Synth*, an implementation of our synthetic reference image approach to correcting EPI distortion. We begin by describing the mathematical framework we use to generate synthetic BOLD images as well as an approach for using them to correct distortion. Then, we demonstrate that undistorted synthetic images constructed using this framework are quantitatively more similar to real BOLD images than T1w or T2w images. Building on these results, we compare this approach to distortion correction against other commonly used MRI field map-based and field map-less correction procedures. We performed these comparisons in a subset of the Adolescent Brain Cognitive Development (ABCD) study dataset [24] in order to assess performance across a variety of scanners, sites and brain geometries [25]. Additionally, we compared performance in the Midnight Scan Club (MSC) dataset, which consists of ten highly sampled individuals [26]. The repeated sampling of MSC participants allowed us to assess the reliability of various methods of EPI distortion correction when applied to the same participants across multiple acquisitions. In order to facilitate the use and development of this approach by others, we provide a software package called *Synth. Synth* can be used to correct fMRI distortions using a synthetic image as an alignment target. The *Synth* software may be used to augment existing fMRI preprocessing pipelines, or explored by researchers interested in incorporating variations on these themes into their MRI registration procedures.

## 2. Methods

### 2.1. Creating an undistorted synthetic BOLD image using Synth

Our approach to correcting EPI distortion is first to create a synthetic BOLD image based on a participant’s undistorted T1w and T2w images, and then use this synthetic image as a reference for nonlinearly aligning a participants’ real BOLD image. Here, we refer to any real image that *Synth* attempts to synthesize as a target image. The ideal synthetic BOLD image will have three properties: 1) it will match the real BOLD target image in terms of overall signal intensity; 2) it will exhibit similar contrast between fluids and tissue types that closely correspond to what is observed in the BOLD target image and; 3) It will account for the difference in spatial resolution between typically high-resolution anatomical images and typically low-resolution BOLD images. Thus, the synthetic image can be represented by the following model:

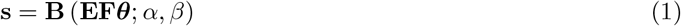

The left hand side of Eqn. 1, **s**, represents a final synthetic image. Working from inside to outside, the right hand side of Eqn. 1 represents the following: the matrix, **F**, is the decomposition of pre-aligned T1w and T2w anatomical images into a set of basis vectors (e.g., radial basis functions — as in Figure 1a, or b-splines, etc.) in order to model the continuous relationship between the voxel intensities of the anatomical images and BOLD target image; ***θ*** comprises the weights for each column of **F. E** is a blurring operator modeling the effective difference in spatial resolution between the anatomical images and BOLD target image. In our implementation of *Synth*, the blurring operator, **E**, is effected with an Epanachnikov smoothing kernel, chosen for its optimal noise-resolution properties [27]. Finally, **B** is a global image contrast operator modeled as a cumulative beta distribution, parameterized by scalars *α* and *β*. When ***θ***, *α*, and *β* are at the optimum solution, **s** represents a synthetic image with the geometry of the undistorted source images and the contrast properties of the BOLD target image. Solving for the optimal ***θ***, *α*, and *β* requires solving a joint optimization problem, which is discussed in the next section.

**Figure 1:**
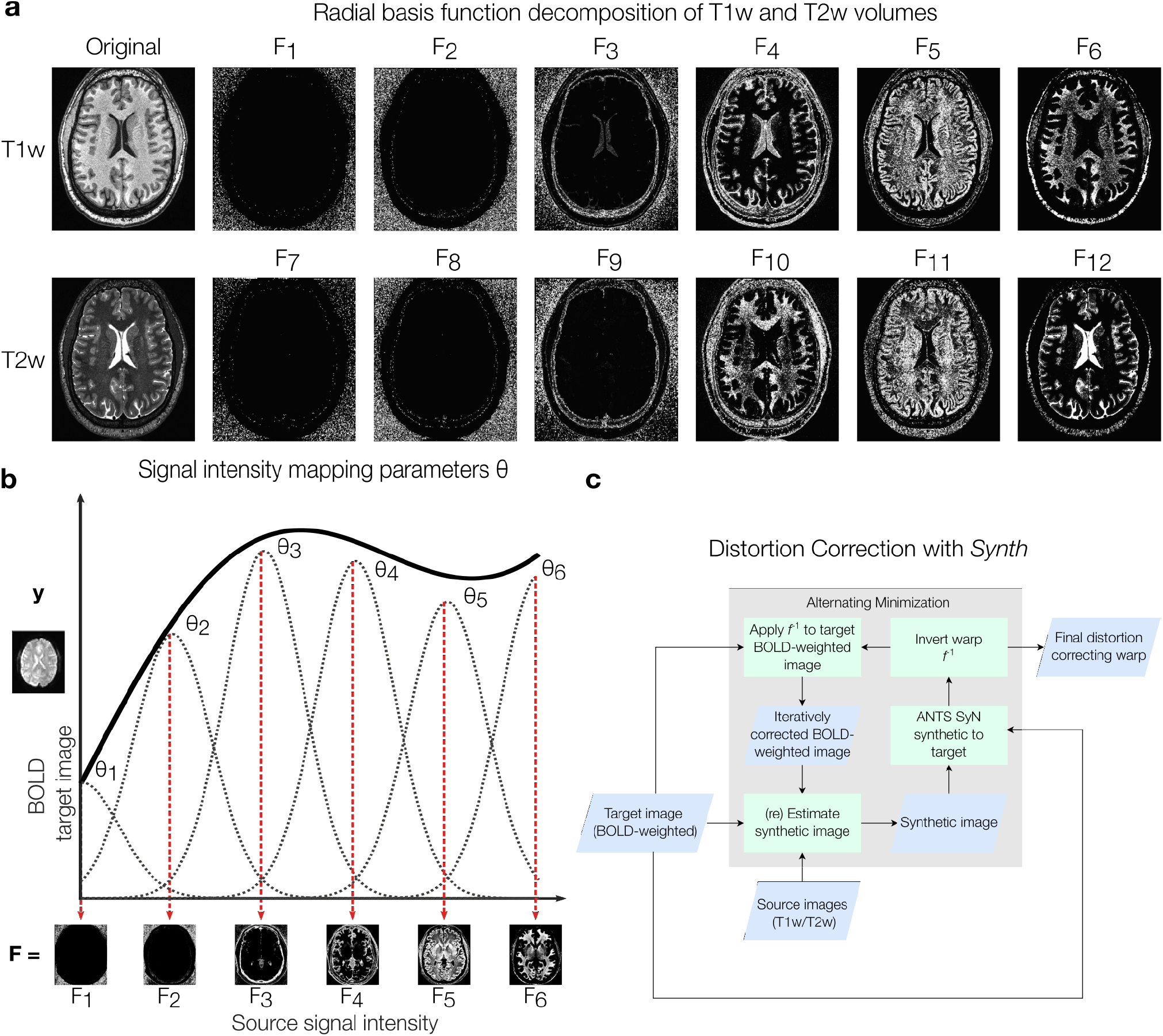
*Synth* synthetic image parameters. **(a)** Illustrative radial basis function (RBF) decomposition of T1w and T2w source images that comprise the columns of **F** (Eqn. 1). RBF decomposition divides source images into smooth ‘bins’ of signal intensity so that the mapping between source and target image intensities can be estimated. Voxel intensities of a target image, e.g., a BOLD image can be modeled as linear combinations of T1w and T2w RBF images. For visualization purposes, we depict a six component decomposition of the T1w image only; *Synth* allows for an arbitrary degree of image decomposition to maximize flexibility in modeling target images with different contrast properties. **(b)** A portion of a hypothetical nonlinear relationship between voxel intensity values observed between source image RBF components (e.g., T2w; x-axis) and voxel intensity values observed in a target image (e.g., BOLD). This image depicts the **F*θ*** portion of Eqn. 1. **(c)** Overview visual of the *Synth* algorithm. The displacement field, *f*, that aligns the initial synthetic image to the target image is estimated using the SyN algorithm. The resulting displacement field, *f*, is inverted to produce the distortion correcting warp, *f* ^−1^, which reduces the target BOLD image distortion. This improves the correspondence between the target and synthetic images, and by proxy, the source images, allowing for improved estimates of a new synthetic image. The distortion correcting warp, *f* ^−1^, is updated in an alternating minimization scheme, during which the synthetic image is refined after each improved estimate of *f* ^−1^.

The columns of the matrix, **F**, are composed of a radial basis function (RBF) decomposition of the source T1w and T2w anatomical images. This decomposition is visualized in Figure 1a, where the T1w and T2w input images have been divided into smooth voxel intensity ‘bins’, each of which corresponds to an RBF component. Each of these components are weighted by the parameter, ***θ***, such that the weighted sum of these components replicates the contrast properties of the BOLD target image as visualized in Figure 1b. *Synth* allows for a user-defined number of intensity ‘bins’ into which the T1w/T2w images are decomposed. The choice of number of intensity bins is dictated by a tradeoff between computational complexity and the accuracy of the synthetic image. As the number of RBF components in **F** increases, so too do the memory requirements. For the synthetic images in the presented results, a 24 component RBF decomposition of both T1w/T2w images (12 components of T1w; 12 components of T2w) at 1 mm isotropic resolution consumed ∼ 20 GB of RAM.

### 2.2. Estimation of synthetic image and distortion-correction warp

To relate the synthetic BOLD image to the real BOLD target image, we model the distortion of the synthetic image as a non-linear warp, *f*, constrained to operate in the phase-encoding direction. This model is described by:

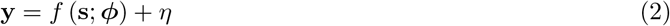

Here, **y** represents the target image (e.g., a distorted BOLD image), ***ϕ*** represents the underlying parameterization of the nonlinear transformation, *f*, and *η* represents additive gaussian noise. By combining Eqn. 1 and Eqn. 2, and solving for the parameters that maximize the correlation coefficient between the real and distorted synthetic image, we obtain the following joint optimization problem:

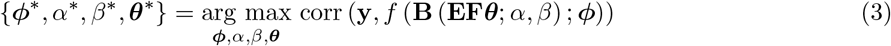

To solve Eqn. 3, we employ an alternating minimization approach, where one parameter is optimized while the others are held fixed over each iteration. ***ϕ*** is initialized with a rigid-body transform so that both synthetic and real BOLD images are closely registered. Once an updated value for ***ϕ*** is found through non-linear registration, the inverse of the estimated nonlinear warp, *f* ^−1^, is then computed. *f* ^−1^ is analogous to a traditional field map correction and maps voxels in a distorted BOLD image, **y**, back to their correct locations. Using the corrected target image, *f* ^−1^ (**y**), an updated value for ***θ*** can be found by solving the linear system *f* ^−1^ (**y**) = **EF*θ***. This updated ***θ*** is used to generate an updated intermediary synthetic image. Finally, the parameters, *α* and *β*, that control the global contrast for the intermediary synthetic image, are computed by minimizing the least squares difference between the synthetic image and distortion corrected BOLD image. The resulting synthetic image is then used for another iteration of non-linear registration.

In regions of the brain with significant distortion and signal dropout affecting the EPI image, there exists no valid mapping between a participant’s anatomical and BOLD images. Therefore *Synth* also allows for the inclusion of a weight volume to reduce the contributions of these when estimating the parameters, ***θ***, *α* and *β*. For the presented results, we down-weighted the contributions of voxels in high-distortion areas (Supplemental Figure 3).

With the application of *f* ^−1^ to the real BOLD data during each iteration, geometric correspondences between anatomical and functional images are improved. Because the quality and accuracy of the intermediary synthetic BOLD images, depends on how closely registered the anatomical and true BOLD images are, modest improvements to the synthetic image can be achieved after each iteration of the *Synth* algorithm. By default, *Synth* repeats this process for 3 iterations. The full procedure for solving Eqn. 3 is outlined in Algorithm 1 (see Section 6.7 in Supplemental Methods) and Figure 1c.

In principle, many of the pre-existing nonlinear registration utilities are suitable for estimating *f* and *f* ^−1^, owing either to their methods of diffeomorphic warp construction which guarantee the existence of an invertible warp —as is the case with AFNI’s 3dQwarp [28] and ANTs SyN [29]— or their ability to project a potentially non-invertible warp onto a “nearest invertible” space— as is the case with FSL’s FNIRT [30]. For the results presented here and in the reference preprocessing scripts associated with this manuscript, we used the ANTs SyN algorithm with a local cross correlation metric for estimating all non-linear warps (see Section 6.6 in Supplemental Methods). ANTs SyN was chosen for its reliable high performance in non-linear warp estimation [31, 32, 19].

### 2.3. Description of MRI datasets and processing

We assessed the quality of distortion corrections produced by direct mapping to *Synth*-generated synthetic images in two different contexts representing important use cases for fMRI data. First, we examined a subset of 100 participants selected randomly from the Adolescent Brain Cognitive Development (ABCD) study dataset (median age: 9.85 years; min: 9 years; max: 11 years). Our random sample included data acquired using GE, Philips, and Siemens scanners (see Table 1 in Supplemental Material). Second, we evaluated *Synth*’s performance on the Midnight Scan Club (MSC) dataset, which consists of resting state fMRI scans acquired from 10 participants on 10 separate occasions (300 minutes of resting state fMRI data/participant). The MSC precision functional mapping (PFM) [33, 26, 34, 35, 36, 37, 38, 39, 40, 41, 42, 43, 44] dataset allowed us to assess session-to-session reliability of EPI distortion correction schemes across multiple sessions for the same participant. Importantly, image acquisition parameters for ABCD and MSC datasets differ significantly allowing us to assess the performance of *Synth* distortion correction on images with a range of contrast and levels of detail. Image acquisition parameters have been reported in detail elsewhere [45, 26].

**Table 1:** Distribution of datasets acquired by specific scanner models

| Manufacturer | Model | Count |
| --- | --- | --- |
| GE MEDICAL SYSTEMS | DISCOVERY MR750 | 18 |
| GE MEDICAL SYSTEMS | SIGNA Creator | 1 |
| Philips Medical Systems | Achieva dStream | 1 |
| Philips Medical Systems | Ingenia | 6 |
| SIEMENS | Prisma | 33 |
| SIEMENS | Prisma Fit | 41 |

We compared the effectiveness of this approach against five widely used distortion correction algorithms. Three of these approaches rely on separately acquired field map data (FSL *fugue*; FSL *topup*; and AFNI’s 3dQwarp) while the fourth and fifth approaches implement alternative field map-less distortion correction methods, SyN-SDC and Synb0-DisCo. FSL field map corrections were constructed using either *fugue* (used for double echo field maps acquired in the MSC dataset) or *topup* (used for opposite direction phase encoded field maps in the ABCD dataset). In AFNI, the distortion correction is estimated from opposite phase encoded field maps using a “meet in the middle” nonlinear warp estimation implemented in 3dQwarp [28]. This estimated warp is used to correct the image and combined with a separately estimated rigid body transform to align the functional to the anatomical image. Because 3dQwarp does not operate on double echo field map data, we could not assess its performance on the MSC data set. We therefore only compared *Synth*’s performance to FSL_*fugue*_ and SyN SDC, SynB0-DisCo, and rigid body alignment.

For field map-less approaches, we assessed the SyN-SDC field map-less method that corrects distortion by non-linearly aligning the participant’s EPI and anatomical images while constraining allowable warps to regions known to be strongly affected by distortion [18, 46, 19]. Additionally, we evaluated the Synb0-DisCo method, which uses deep learning methods to synthesize a pseudo infinite bandwidth EPI image as an input to FSL’s *topup* to estimate the distortion correcting warp [21]. As a reference, we also computed each alignment quality metric for a dataset in which only rigid body alignment was performed.

Performance of the different distortion correction approaches was assessed in matched datasets, thus all statistical comparisons were performed within a multi-level modeling framework (fitlme, MATLAB). For these analyses, the metrics produced by the *Synth*-based registration pipeline were modeled as the baseline and differences in performance metrics associated with other approaches were modeled with main effect factors. To account for participant-specific variability, independent of the registration approach, participant identity was modeled as a random effect.

### 2.4. Evaluation Metrics

No consensus exists as to what global metric best quantifies alignment quality. Many metrics used to measure alignment quality tend to bear some relation to the cost functions that are used during registration and therefore directly introduce a risk of confounding circularity. Thus, we computed several metrics comparing the similarity of different features of the field map corrected BOLD images to their associated anatomical images.

#### 2.4.1. Contrast Similarity

To quantify the overall similarity between two aligned images from different modalities, we computed a contrast similarity metric which was defined as the linear correlation coefficient between vectorized versions of corresponding regions of the two images. Correlations were constrained to values that reside within a full brain binary mask.

#### 2.4.2. Normalized mutual information (NMI)

A quantification of global image similarity determined by the amount of information shared between the time-averaged BOLD image from each session and its associated T1w/T2w anatomical images (e.g., T1w-BOLD NMI, T2w-BOLD NMI).

#### 2.4.3. Edge alignment

A quantification of alignment between any high-contrast edges existing in two images. It is defined as the correlation coefficient between the gradient magnitude images of the time-averaged BOLD image and T1w/T2w images within a whole brain mask.

#### 2.4.4. Segmentation alignment

In BOLD images, gray matter, white matter, and CSF tend to exhibit distinct intensity values. When anatomical and BOLD images are well aligned, segmentation maps derived from anatomical images and overlaid on the BOLD image should correspond well to the tissue types of the BOLD image. In well aligned images, the distributions of intensity values of BOLD image voxels for a particular tissue type should tend to be more distinct and separable than for poorly aligned images. Here, separability is defined as the ability of a linear classifier (i.e. thresholding) to correctly distinguish between two tissue types. Each segmentation metric represents the area under the curve (AUC) of a receiver operating characteristic curve (ROC) generated from distributions of voxels delineated by two segmented tissue types. Each metric differs on the two tissue types selected. The segmentation of each tissue is limited to voxels ‘adjacent’ to the other tissue - where ‘adjacent’ refers to the set of all voxels in the 1st tissue that shares a face with a voxel in the 2nd tissue and vice-versa. AUC_*gw*_ represents the separability of adjacent gray and white matter voxels, AUC_*ie*_ represents the separability of adjacent voxels between the interior and exterior of the brain, and AUC_*vw*_ represents the separability of adjacent voxels in the ventricular cerebrospinal fluid and white matter.

#### 2.4.5. Local alignment

Often, approaches to image registration rely on minimizing or maximizing a global distance or similarity metric (e.g., least-squares or mutual information) computed across an entire image volume. These metrics can be strongly influenced by the presence of intensity bias fields, which are low spatial frequency modulation of image intensity often caused by suboptimal participant placement or poor shimming. Additionally, experimental evidence has suggested that inaccurate registration can, in some situations, produce high values for many global measures of image alignment [15]. We therefore sought to compare the performance of each approach to EPI distortion correction with respect to a measure of local image similarity that would be less sensitive to these influences.

To do this, we implemented a ‘spotlight’ analysis examining 7×7×7 voxel regions (3 mm isotropic voxels) of the BOLD images produced by each method and quantified its similarity to the corresponding region of the participant’s T1w and T2w images with an *R*^2^ metric. Image registration quality near gray matter is typically of greatest interest, hence we summarized local T1w-BOLD and T2w-BOLD similarity by computing the average spotlight *R*^2^ across all gray matter voxels, as labeled by the participant’s Freesurfer segmentation (freesurfer citation).

#### 2.4.6. Intra-participant stability

We used the MSC dataset to investigate how different distortion correction procedures affect metrics of inter-session stability for fMRI data. We reasoned that improving EPI image consistency would also reduce the variability of the intensities of individual voxels across sessions. To test this, we constructed time-average BOLD images for each session contributed by a participant. We temporally concatenated each of these images, forming a pseudo-time series reflecting session-to-session changes in the BOLD images. Then, we computed a metric, session-signal-to-noise (s-SNR, analogous to traditional t-SNR), for each voxel defined as the mean intensity of a voxel, *μ*, divided by the standard deviation of its intensity across sessions, *σ*. Lastly, we summarized alignment stability for each participant by averaging s-SNR across all voxels residing within a whole brain mask defined by their participant-specific FreeSurfer parcellation.

Operating on the assumption that less reliable field map correction procedures would introduce additional session-to-session variability in resting state functional connectivity (RSFC) matrices (pairwise correlations between voxel time series), we measured the similarity of each pair of RSFC matrices contributed by a given participant. We identically preprocessed (see Section 6.1.3 in Supplemental Methods) the MSC datasets produced by each registration pipeline and computed the RSFC BOLD-signal correlation matrices for all gray matter voxels. For each session, we extracted the upper triangle of the associated correlation matrix. Finally, we quantified resting state correlation matrix stability by computing the pairwise correlation between the upper triangles of each unique pair of a participant’s resting state matrices.

## 3. Results

### 3.1. Synth images consistently match the contrast of BOLD images

We reasoned that anatomically-based reference images with greater contrast similarity to BOLD images would serve as a more reliable reference for distortion correction than images with lower contrast similarity. To assess differences in contrast similarity between BOLD and T1w, T2w and synthetic images, we calculated the linear correlation coefficient between participants’ BOLD images and each of their T1w, T2w and synthetic images within a whole brain mask. By default, *Synth* uses a model that includes a 12-component radial basis function (RBF) decomposition of the T1w/T2w source images along with pair-wise T1w/T2w interaction terms to create synthetic BOLD images (Figure 1). This model was sufficient to produce synthetic images with comparable BOLD contrast similarity in both ABCD and MSC datasets (Figure 2a,b). In both datasets, synthetic images exhibited significantly increased contrast similarity to BOLD images compared to T1w and T2w images (Figure 2c). Contrast similarity between the BOLD and *Synth* images was consistently highest. Mixed-effects model comparisons in the ABCD data set revealed that the differences between *Synth* contrast similarity and T1w/T2w contrast similarity were both significant: (T1w-*Synth*= −0.424; *p* < 0.001; *t* = −46.18; *df* = 297) and (T2w-*Synth*= −0.18; *p* < 0.001; *t* = −19.62; *df* = 297). The same pattern held in the MSC dataset: (T1w-*Synth*= −0.374; *p* < 0.001; *t* = −49.64; *df* = 297) and (T2w-*Synth*= −0.274; *p* < 0.001; *t* = −36.4; *df* = 297).

**Figure 2:**
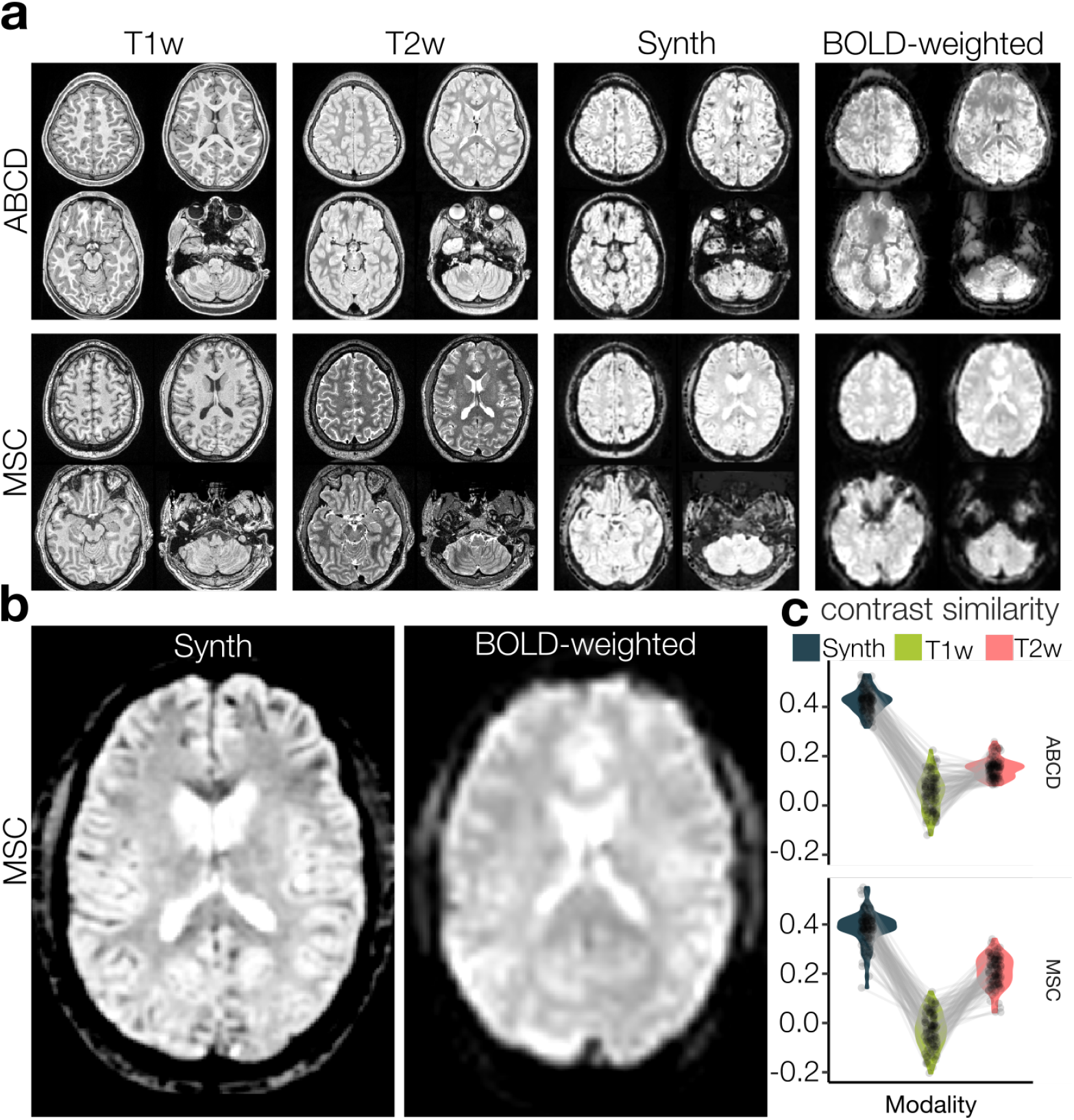
Image contrast similarity between T1w, T2w, *Synth* and BOLD images. **(a)** T1w, T2w, synthetic, and corresponding affine aligned BOLD images (sample participants; ABCD top, MSC bottom). **(b)** Enlarged view of an axial slice of the Synthetic (left) and real BOLD (right) images (MSC example). **(c)** Average contrast similarity between BOLD images and associated anatomical and *Synth* images across 100 participants from the ABCD dataset (upper). Average contrast similarity between a true BOLD image and associated anatomical and synthetic functional images for each participant in the MSC dataset (lower; 10 scans per participant).

### 3.2. Synth outperforms existing field map-less methods on global measures of image alignment

We assessed *Synth*’s distortion correction performance (Figure 3) against existing field map-based (AFNI 3dQWarp; FSL *topup*; FSL *fugue*) and field map-less (SyN-SDC; SynB0-DisCo; Rigid-body) distortion correction methods using both established and novel metrics that quantify global image similarity between the BOLD image and the associated T1w/T2w anatomical images (Figure 4). These metrics included normalized mutual information; the correlation coefficient between the image gradients; and segmentation alignment. Here, we outline *Synth* registration performance for each metric described above as compared to the highest performing competitor. *Synth* registration tended to produce BOLD images with the highest registration quality metrics for both ABCD and MSC data. Comparisons are reported in the form of statistical contrasts corresponding to the mean difference between registration metrics (e.g., Method A-Method B). Full statistical tables comparing all methods are included in Table 1 of Supplemental Material. Illustrative comparisons between FSL fugue/topup and Synth are included in Supplemental Figures 1 and 2.

**Figure 3:**
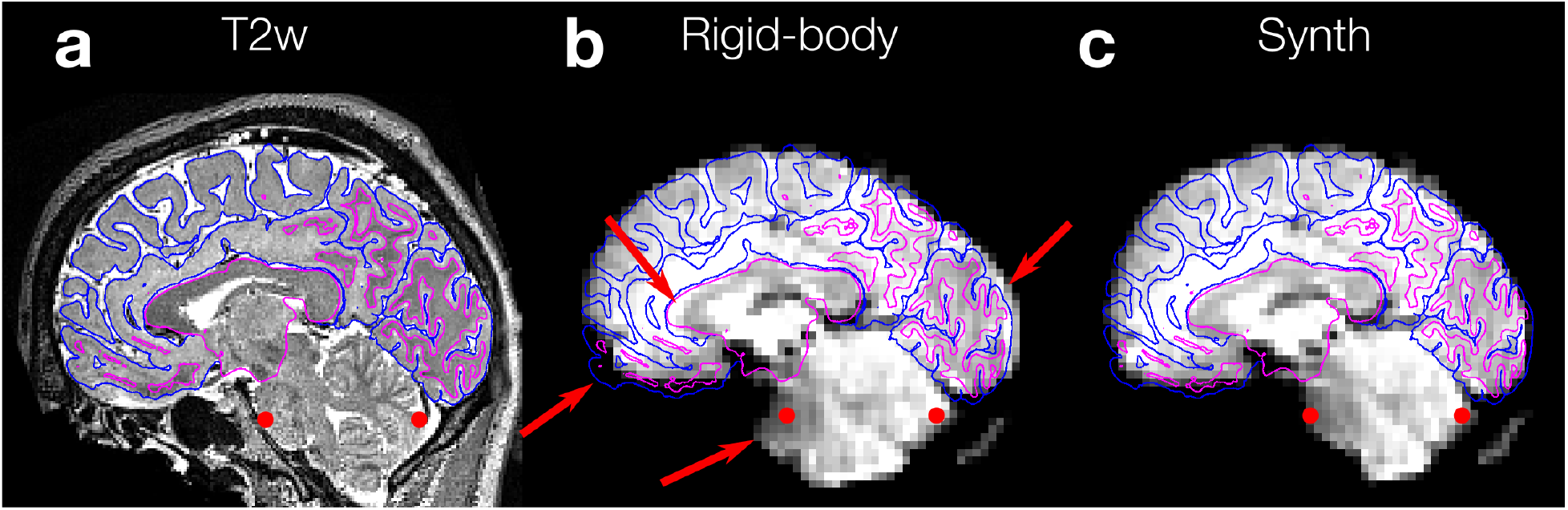
Example of distortion correction with *Synth* compared to rigid-body alignment. **(a)** Parasagittal slice of T2w image from an example MSC participant. Fuschia and blue lines indicate gray/white matter boundary estimated by freesurfer segmentation of the associated T1w image. **(b)** Corresponding slice of functional EPI image aligned to the anatomical image using rigid body alignment procedure implemented in FSL’s FLIRT. Red arrows indicate regions of local misalignment due to image distortion. Red dots indicate anatomical fiducial markers to aid in assessing distortion correction applied to cerebellum and pons. **(c)** Identical slice through EPI image corrected using *Synth*. Note improved local alignment of corpus callosum, occipital pole, pons, and orbitofrontal cortex.

**Figure 4:**
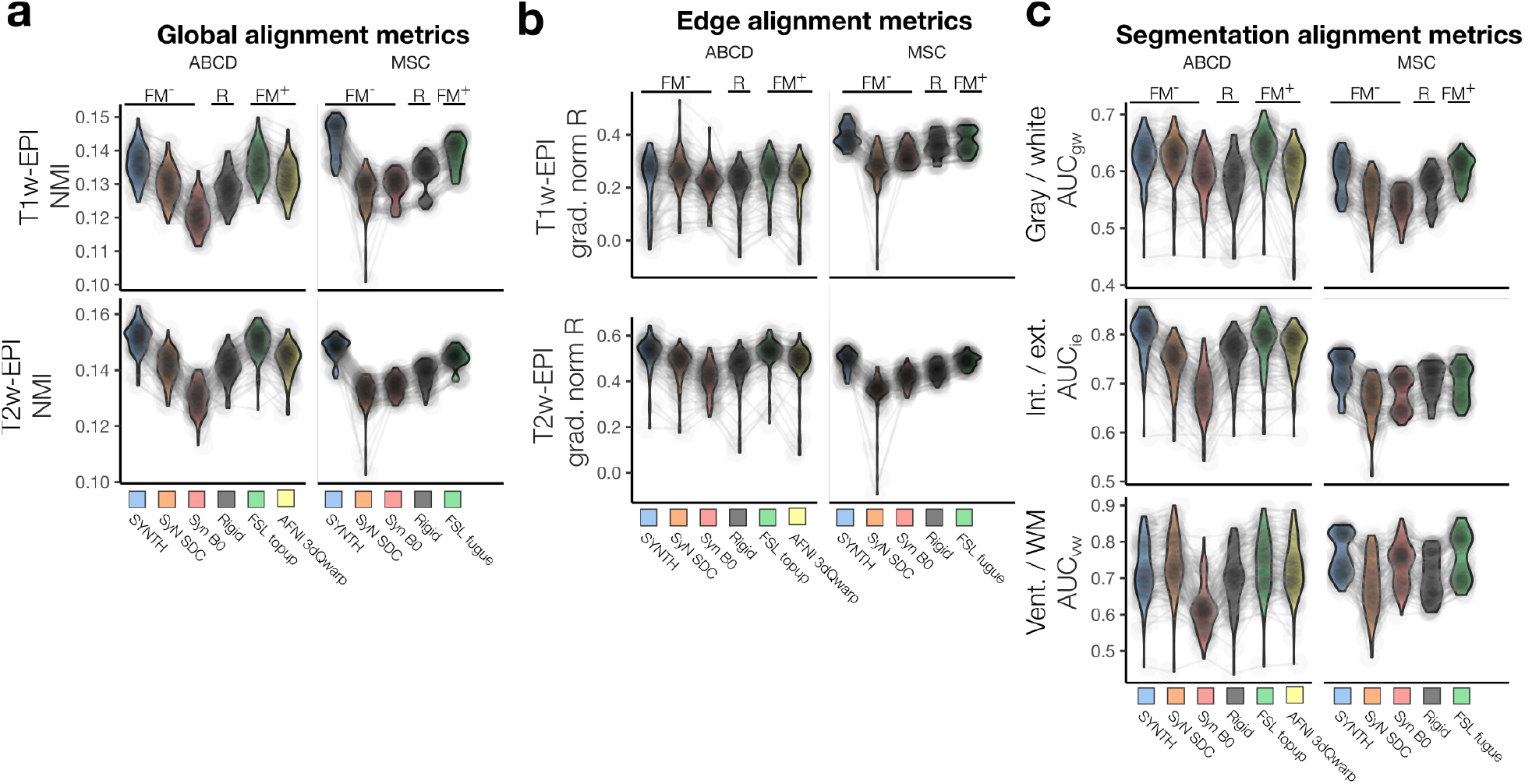
Alignment metric comparisons for distortion correction pipelines. Violin plots depict the distributions of each alignment metric for ABCD (left column) and MSC (right column) datasets. Horizontal bars group assessment metrics for field map-less distortion correction methods (FM-), rigid body (R) and field map based (FM+) distortion correction methods. **(a)** Global similarity: Normalized mutual information (NMI) shared between the registered BOLD images and their associated T1w/T2w anatomical images to assess global similarity. **(b)** Edge alignment: Correlation coefficient between the gradient magnitude images for BOLD and T1w/T2w images for a metric of edge alignment. **(c)** Segmentation alignment: Metrics that quantify the separability of anatomical features in BOLD data base on the Freesurfer segmentation of their anatomical T1w images.

First, we examined a well-established image similarity metric, normalized mutual information (NMI). In the ABCD dataset, NMI between BOLD and T1w images was not significantly different (FSL_*topup*_-*Synth*= −4e-5; *p* = 0.9; *t* = −0.13; *df* = 594). In contrast, NMI for T2w images was significantly higher for *Synth* images than those aligned by FSL_*topup*_ (FSL_*topup*_-*Synth*= − 1.7e-3; *p* < 0.001; *t* = − 3.6; *df* = 594). *Synth* alignment produced BOLD images with the highest NMI shared between T1w/T2w images in the MSC dataset as well (FSL_*fugue*_-*Synth*= − 4e-3; *p* < 0.001; *t* = − 12.4; *df* = 495) and (FSL_*fugue*_-*Synth*= − 3.5e-3; *p* < 0.001; *t* = − 10.1; *df* = 495) (Figure 4a).

Next, we examined the quality of alignments emphasizing registration of high contrast boundaries. BOLD-T1w edge alignments produced by *Synth* were outperformed by those produced by SyN SDC (SyN SDC-*Synth*= 3.1e-2; *p* < 0.001; *t* = 5.23; *df* = 594) in the ABCD dataset; for T2w gradient magnitude images, there was no significant difference between *Synth* and the highest performing competitor, FSL_*topup*_ (FSL_*topup*_-*Synth*= −4.6e-3; *p* = 0.39; *t* = −0.86; *df* = 594). In the MSC dataset, its closest competitor varied by modality. For BOLD-T1w edge alignment, FSL_*fugue*_ was the highest performing competitor, but still produced edge alignments that were significantly lower than those produced by *Synth* distortion correction (FSL_*fugue*_-*Synth*= −0.023; *p* < 0.001; *t* = −4.8; *df* = 495). For T2w images in the MSC dataset, FSL_*fugue*_ was once again the closest competitor but did not perform significantly better than *Synth*, (FSL_*fugue*_-*Synth*= 7.8e-5; *p* = 0.88; *t* = 0.15; *df* = 495) (Figure 4b).

In addition, we examined the quality of the alignment of BOLD images to anatomical images segmented by tissue type (*gw*: gray/white matter; *vw*: ventricles/white matter; *ie*: interior/exterior of brain) based on the participants’ freesurfer segmentation. These metrics quantify the separability of BOLD image voxel intensities based on anatomically defined segmentation (higher AUC [area under the receiver operating characteristic curve] is better). AUC_*gw*_ image segmentation alignment metrics for *Synth* were significantly lower than those produced by FSL_*topup*_ in the ABCD dataset (FSL_*topup*_-*Synth*= 1.6e-2; *p* < 0.001; *t* = 5.8; *df* = 594) and MSC (FSL_*topup*_-*Synth*= 8e-3; *p* < 0.001; *t* = 3.46; *df* = 495) datasets. *Synth*-aligned images exhibited greater AUC_*ie*_ in both ABCD (FSL_*topup*_-*Synth*= −0.016; *p* < 0.001; *t* = −6.11; *df* = 594) and MSC (Rigid-*Synth*= −1.9e-2; *p* < 0.001; *t* = −8.4; *df* = 495) datasets. We also found that the SyN SDC pipeline produced the greatest AUC_*vw*_ values (SyN SDC-*Synth*= 0.021; *p* < 0.001; *t* = 3.65; *df* = 594) for the ABCD dataset. However, *Synth* performance did not differ significantly from the highest performing competitor, FSL_*fugue*_ (FSL_*fugue*_-*Synth*= −4.4e-3; *p* = 0.34; *t* = −1.0; *df* = 495) in the MSC dataset (Figure 4c).

### 3.3. Synth improves local measures of anatomical and functional image alignment

Existing experimental evidence indicates that field map correction quality varies regionally and depends on the method used to estimate the B0 inhomogeneity [10]. Consequently, to identify whether *Synth* was better at correcting certain brain regions, we carried out a winner-take-all (WTA) analysis across all voxels by determining which method produced the greatest average local similarity between T1w and T2w anatomical images (*R*^2^ computed within a 7×7×7 voxel spotlight). Each distortion correction method exhibited regions of the brain for which they performed consistently better than other distortion correction methods. Although the structure of the WTA maps varied between datasets, they shared some consistent features (Figure 5).

**Figure 5:**
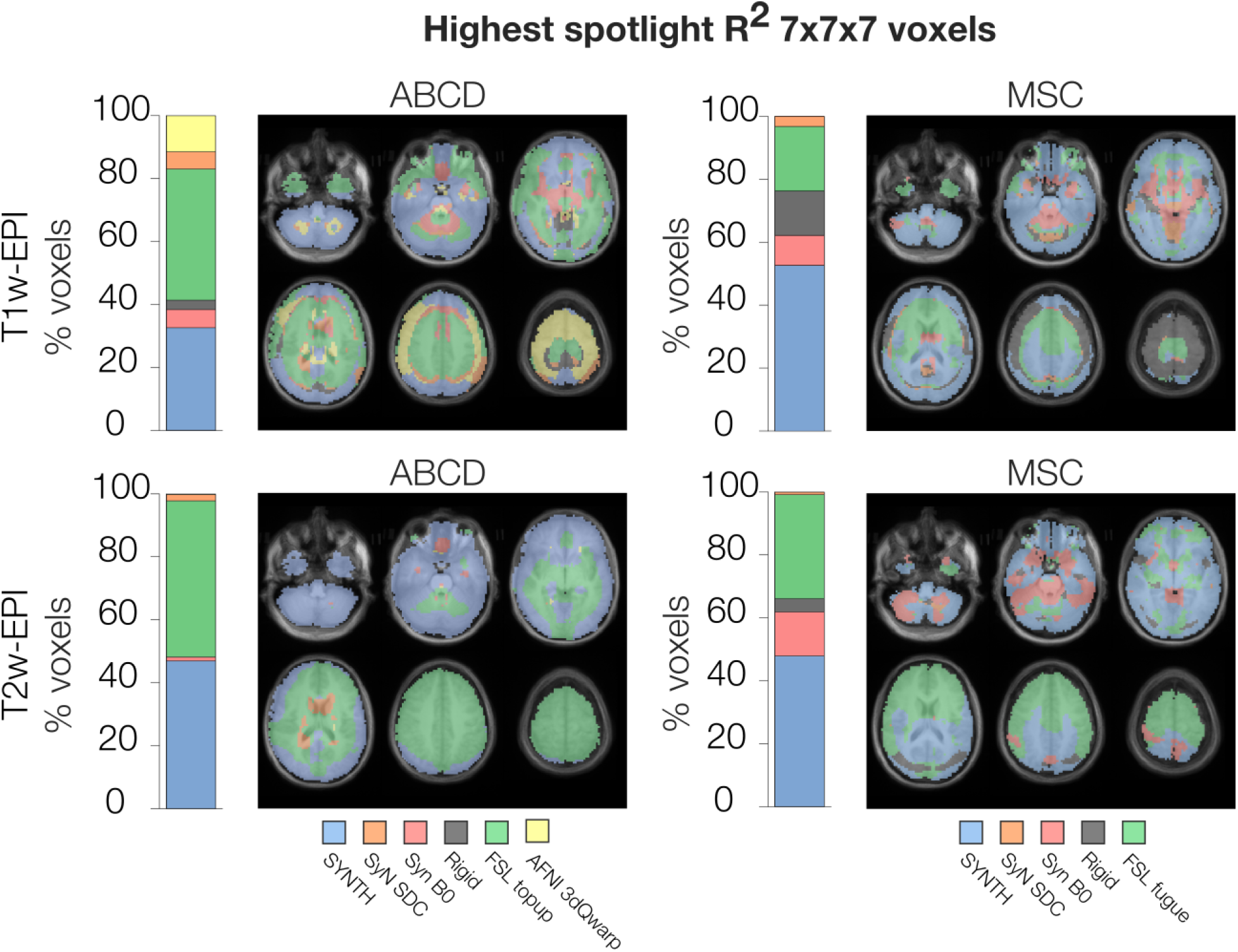
Regional variation in distortion correction performance. Summary of local similarity between the BOLD images produced by each registration pipeline and their associated T1w and T2w anatomical images. Winner-take-all maps depicting, for each voxel, the pipeline that produced the greatest average local correlation between corresponding regions of BOLD and T1w/T2w images. The color of each voxel depicts which registration pipeline produced, on average, the greatest spatial R2 values computed over a centered 7×7×7 cube. Bars to the left of each set of images depict the percentage of voxels “won” by each registration pipeline.

In both datasets, *Synth* produced either the largest, or second largest regions of high quality local alignment, regardless of anatomical image modality (i.e., T1w, T2w). In the MSC data set, *Synth* generally had the best regional correction performance across the entire brain. For the ABCD dataset, the best regional distortion correction performance was split between FSL_*topup*_ and *Synth*, where FSL_*topup*_ provided better local alignment to T1w images over a large swathe of the superior frontal cortex while *Synth*’s particular strengths were in improving local registration of cerebellar, brainstem, orbitofrontal and occipital regions with respect to a participant’s T2w image. Overall, FSL_*topup*_ won the greatest percentage of voxels in the T1w and T2w comparisons for the ABCD dataset, (T1w: FSL_*topup*_= 42%; *Synth*= 33%; T2w: FSL_*topup*_= 50%; *Synth*= 47%). While *Synth* won the greatest percentage of voxels in the T1w and T2w comparisons over FSLfugue (T1w: FSL_*fugue*_= 21%; *Synth*= 53%; T2w: FSL_*fugue*_= 33%; *Synth*= 48%) for the MSC dataset. This indicates that *Synth* can perform at parity with state-of-the-art field map-based approaches.

The analysis of functional images depends to a greater extent on the quality of gray matter alignment and less so on the alignment quality of other voxels representing other tissue types. Accordingly, we summarized local gray matter alignment performance of each distortion correction method by computing the average spotlight *R*^2^ across only the gray matter voxels. We observed that *Synth* and FSL_*topup*_ registration pipelines produced similar quality of local registration to the participants’ T1w image (FSL_*topup*_-*Synth*= −0.002; *p* = 0.11; *t* = −1.6; *df* = 594), in the ABCD dataset. *Synth* performance did not differ significantly from the highest performing competitor, FSL_*topup*_, for average local similarity between BOLD and T2w images (FSL_*topup*_-*Synth*= −5.4e-3; *p* = 0.081; *t* = −1.75; *df* = 594). *Synth* registration produced BOLD images with the greatest average local similarity for associated T1w images (FSL_*fugue*_-*Synth*= −1.8e-2; *p* < 0.001; *t* = −9.2; *df* = 495) and did not differ significantly from the highest performing competitor, FSL_*fugue*_ for T2w images (FSL_*fugue*_-*Synth*= −9e-4; *p* = 0.67; *t* = −0.42; *df* = 495) in the MSC dataset (Figure 6).

**Figure 6:**
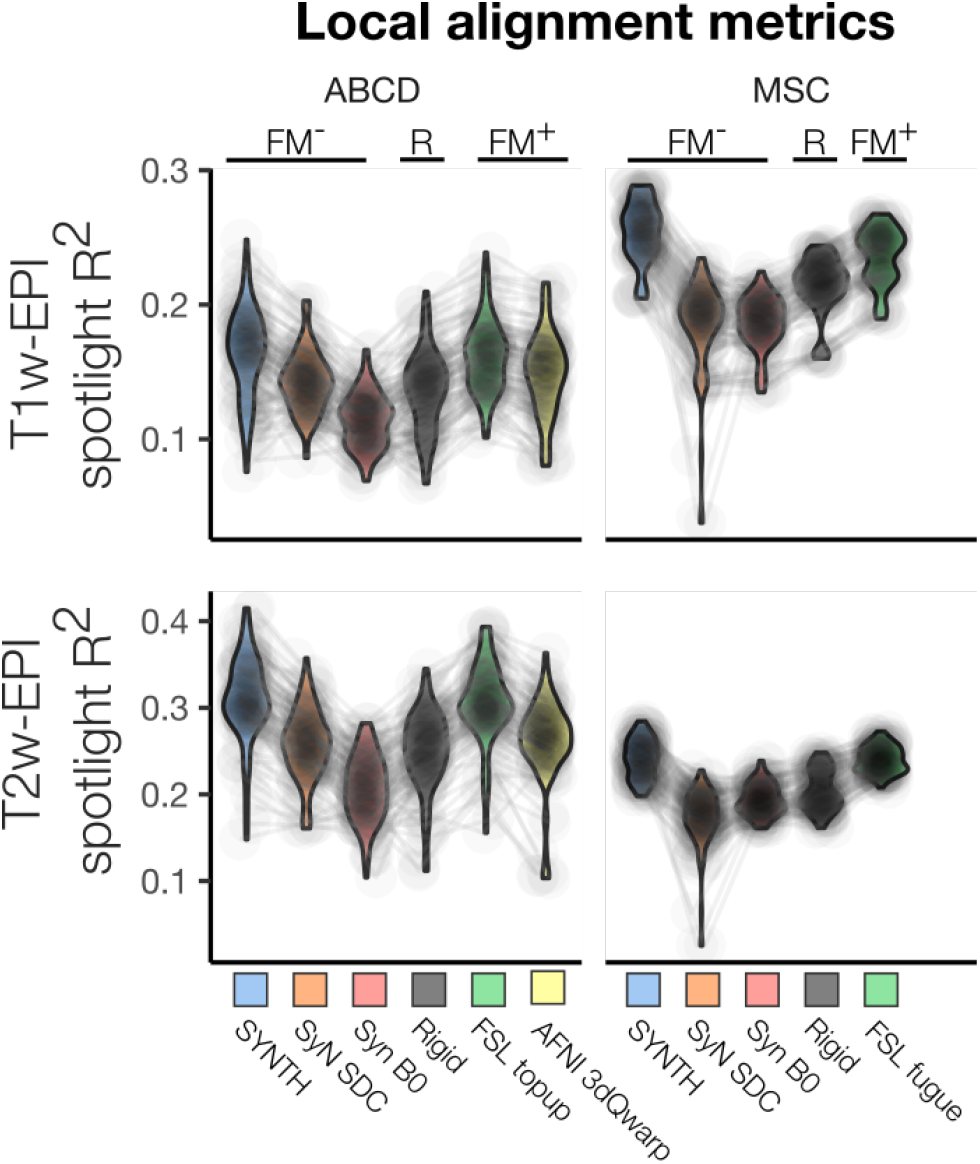
Average local image similarity across all gray matter voxels. Summary of local similarity between the BOLD-weighted images produced by each registration pipeline and their associated T1w/T2w anatomical images. Each plot depicts the distributions of average spotlight R2 value between each participant’s BOLD image and T1w/T2w images computed across all gray matter voxels.

### 3.4. Synth improves intra-participant consistency across sessions

The primary goal of applying distortion correction to EPI images is to improve data quality by reducing variability caused by poor alignment. It is therefore important to know the extent to which a particular approach to distortion correction introduces additional measurement variability. By applying different distortion correction strategies to the repeated-measures in the MSC dataset, we assessed how each method contributed to within-participant structural and functional variability.

First, we examined each voxel’s average intensity across all sessions normalized by its standard deviation across sessions. We term this stability metric, session-to-session SNR (s-SNR). We observed that for *Synth*-aligned images, s-SNR was not significantly different than that produced by the highest performing competitor, FSL_*fugue*_ (FSL_*fugue*_-*Synth*= 0.79; *p* = 0.52; *t* = 0.6; *df* = 45) (Figure 7a).

**Figure 7:**
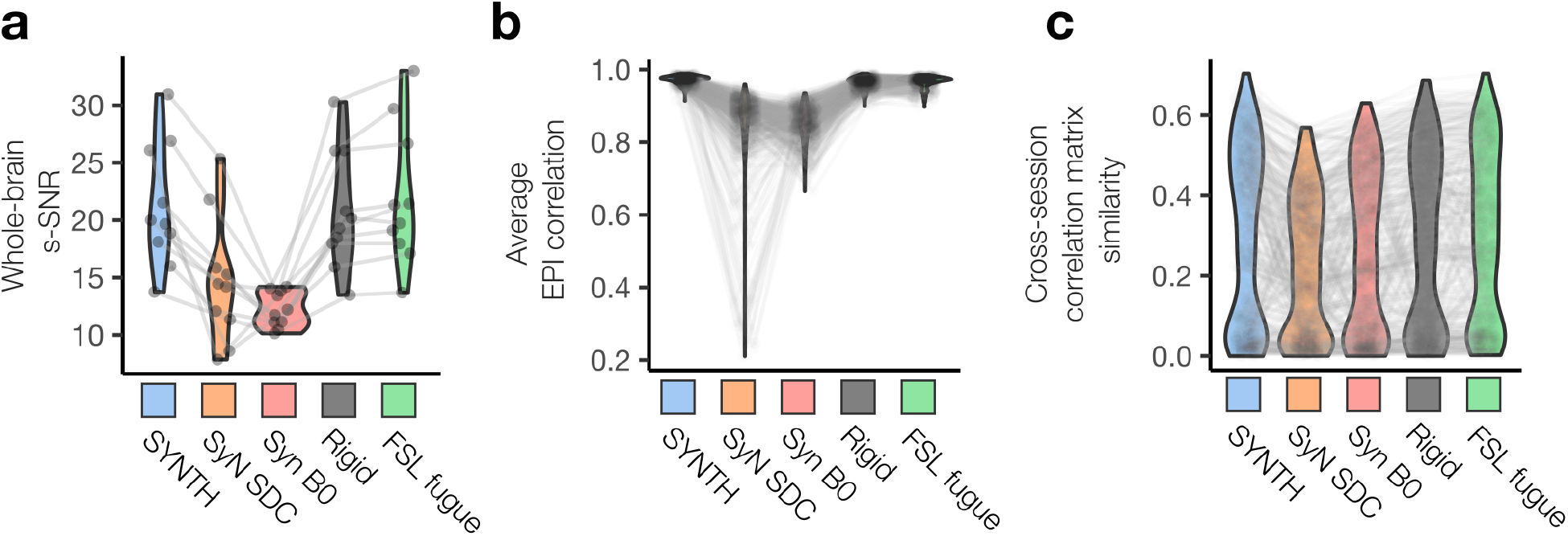
Cross-session structural and functional reliability metrics. Repeated measurements of the same 10 individuals in the MSC dataset allows us to assess the reliability of different approaches to EPI distortion correction. **(a)** Average voxelwise session-SNR (s-SNR) computed within a whole brain mask. s-SNR was computed as 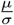, where *μ* is the voxel intensity of the time average BOLD image averaged across all sessions, and *σ* is the standard deviation. **(b)** Distributions of the mean pairwise correlation between time-averaged resting state BOLD volumes. **(c)** Similarity between the BOLD signal correlation matrices from each unique pair of sessions. Each of the 10 participants contributed 10 resting state datasets, corresponding to 45 unique session pairs per participant. Similarity is quantified as the correlation between the vectorized upper triangles of the pairs of BOLD signal correlation matrices. Note that these similarity values are smaller than those reported elsewhere for this same dataset as a consequence of the minimal spatial smoothing and the analyses being applied at the voxel as opposed to parcel level.

Next, we quantified global similarity of time averaged BOLD images for each pair of sessions contributed by a participant. We extracted the region of the aligned time-averaged BOLD volumes containing just brain matter (as classified by the FreeSurfer segmentation of the participant’s anatomical images) from each session. Then, for each participant, we computed the average linear correlation coefficient across all pairs of sessions. Owing to the general stability of a given participant’s brain geometry on the time scales over which the MSC dataset was collected, inter-session linear correlation coefficients between scan sessions tended to be quite high across all methods (0.94 − 0.99). While *Synth* produced BOLD images with the greatest session-to-session similarity, it did not produce images that were significantly more stable than those produced by rigid body alignment alone (Rigid-*Synth*= −7.4e-3; *p* = 0.28; *t* = 1.8; *df* = 2245) (Figure 7b).

Finally, we examined how the choice of distortion correction approach affects the stability of resting state functional connectivity (RSFC) data. To do this, we created voxel-level RSFC matrices for each session of data and computed RSFC similarity between pairs of sessions as the correlation between upper triangular portions of the RSFC matrices. Mixed effects analysis of similarity between unique session pairs revealed RSFC correlation matrices from *Synth* aligned datasets were not significantly less stable than those produced by the highest performing competitor, FSL_*fugue*_ (FSL_*fugue*_-*Synth*= 4.6e-3; *p* = 0.5; *t* = 0.67; *df* = 2245) (Figure 7c).

## 4. Discussion

Distorted fMRI images are an inevitable consequence of the trade-offs made to achieve a sampling rate sufficient to measure neurally meaningful fMRI signal variability in Cartesian sampled EPI [47]. Complications in registering distorted functional BOLD images to their undistorted anatomical image counterparts are a perennial challenge faced by neuroimaging researchers. Performed well, proper image registration reduces variability in measurements, improves the statistical power of analyses, and produces significant improvements in localization of responses to experimental manipulations [48, 8, 49, 50]. For these reasons, the quality of EPI distortion correction and anatomical registration is of great concern in every branch of neuroimaging and the primary motivation for developing methods to improve fMRI registration performance.

We hypothesized that the reliability of field map-less approaches could be improved if the underlying contrast properties of the BOLD and anatomical images match more closely. The distortion correction approach that *Synth* employs is to generate a high resolution synthetic functional image that exhibits greater similarity to tissue contrast properties of a BOLD image but is based on the information from the undistorted geometry of a participants’ T1w and T2w anatomical images. The approach implemented in *Synth* for constructing a synthetic BOLD image target can be considered an example of a class of registration procedures termed “mono-modal reduction” [51]. After constructing a synthetic target image, a distortion-correcting warp is constructed in the same way as other direct mapping approaches: by directly mapping to an undistorted target image. Here, we have demonstrated that when a synthetic image is used as a target, the resulting fMRI alignment quality rivals and in some cases exceeds that produced by standard field map methods.

### 4.1. Synth achieves parity with or exceeds the distortion correction performance of alternative methods on most measures of global alignment

We began our exploration of distortion correction performance by examining the commonly used global image similarity metric, normalized mutual information. We found that *Synth* performed comparably to or better than state of the art field map corrections provided by FSL’s *topup* and *fugue* in both ABCD and MSC data sets. One of the challenges to assessing the quality of multi-modal image registration and distortion correction is determining which global metric serves as the optimal summary of final alignment quality. To address this concern, we assessed the performance of each distortion correction approach with several additional global image similarity metrics, each intended to assess alignment with respect to different image features.

The first alternative global similarity metric we examined was the quality of high-contrast edge alignment. We reasoned that these features would be less influenced by residual low spatial frequency bias fields and the differences in tissue contrast properties between images acquired with different modalities (T1w, T2w, BOLD). Here, *Synth*’s performance on cross-modal edge alignment metrics was generally high, either achieving the highest performance or placing in the top two. The single exception to this outcome was observed in the T1w-BOLD edge alignment metrics in the ABCD data set. There, FSL’s *topup* and SyN SDC produced the highest alignment metrics. This result may arise from the fact that the *Synth* pipeline initially registers the BOLD image into the anatomical image space using a T2w reference image, while the FSL and SyN SDC pipelines both register to the anatomical image space using the T1w image as a reference. It may be the case that the choice of image modality (i.e. T1w/T2w) used for the initial BOLD-anatomical alignment biases the edge alignment metric for those two image modalities.

FSL’s *topup* provided the best ventricles and white matter alignment metrics (AUC_*vw*_) in the ABCD dataset followed by the field map-less SyN SDC method. One potential reason for this outcome is that both FSL’s *topup* and SyN SDC pipelines used brain-based registration for their BOLD to anatomical alignments [52]. This registration technique incorporates anatomical segmentation information when guiding anatomical alignments and may improve ventricle/white matter correspondence between the BOLD and anatomical images.

### 4.2. Synth exhibits high performance in regional measures of image alignment

In addition to the challenge of selecting and justifying a particular global metric for measuring multimodal image alignment quality, prior work has shown it does not always provide a sufficient condition for demonstrating optimal alignment [52]. This is because global metrics, while flexible, often cannot fully capture the complicated mapping of voxel intensity values between images acquired with different modalities. While achieving high measures of global image similarity is a necessary condition for quality image alignment, it is important to consider them alongside non-global measures to fully assess alignment [53]. For this reason, we examined small regions of the image (i.e. spotlight), which contain a limited range of tissue types and minimal residual bias field. In this way, we created an alignment metric that assesses image similarity while reducing the influence of non-linearities in the relationship between voxel intensity values of images acquired with different modalities.

Our winner-take-all analyses of local image alignment revealed *Synth* to be among the top performing distortion correction approaches, producing the highest local image quality metrics in large swathes of the cortex, but also performing particularly well in correcting distortions affecting the cerebellum and brain stem. That different distortion correction approaches excelled in correcting different areas of the brain is in line with existing evidence suggesting that the accuracy of standard field map distortion correction methods varies across the brain [10, 54].

One possible reason for *Synth*’s superior performance in the MSC dataset overall, compared to the ABCD dataset, may be due to the dual echo field maps used in the MSC dataset. Prior work has shown that distortion correction with reverse-phase encoding field maps outperform corrections produced by dual echo field maps [54]. Our findings are consistent with these observations, and suggest that *Synth* may provide superior correction to dual-echo field map corrections in datasets in which image acquisition parameters share sufficient similarity to those used for the MSC study [26].

### 4.3. Synth does not introduce session-to-session variability like existing field map-less approaches

We observed that as a general rule, field map-less approaches tended to introduce more session-to-session structural variability than traditional field map-based approaches. *Synth* stood out as a clear exception to this trend by introducing no more session-to-session variability than a simple rigid-body alignment procedure or field map correction. Importantly, because *Synth* distortion correction did not measurably increase the session-to-session variability of RSFC matrices we can conclude that the improvement in BOLD-to-anatomical alignment provided by *Synth* does not come at a hidden cost of increasing the variability of the underlying functional data.

### 4.4. Accounting for differing performance of existing field map-less approaches

*Synth* tended to produce higher quality and more reliable corrections than existing field map-less approaches (SyN SDC and SynB0-DisCo) with which it shares many conceptual similarities. This prompts us to consider the potential reasons why *Synth* performs with greater reliability and accuracy.

In the case of SyN SDC, one reason for decreased reliability may be the use of a group average field map template in order to constrain its solutions. Such a template may not allow sufficient flexibility to align high spatial resolution participant-specific features. SyN SDC also relies on an intermediate atlas alignment stage, which may be a source of additional variability. Because participant motion can influence the structure of the EPI distortion, the use of a field map template that does not account for this effect may reduce overall efficacy.

Both *Synth* and SyN SDC employ a direct mapping approach to correct EPI distortion. The fundamental difference in their approaches is that SyN SDC attempts to directly align to a participants’ T1w image - whose contrast properties are very different from the EPI image-while *Synth* uses an intermediate synthetic image whose contrast properties are more similar to the EPI. That *Synth* proved more effective and reliable than the similar approach implemented in SyN SDC would seem to indicate the importance of matching the contrast properties of target and reference images when attempting to use direct mapping approaches for distortion correction.

With SynB0-DisCo, a synthetic pseudo-infinite bandwidth intermediate EPI image facilitates distortion correction using FSL’s *topup*. The corrections, though, were less accurate and less reliable than those produced by *Synth* and SyN SDC. On the surface, it would seem that the use of a contrast matched intermediate image along with FSL’s state of the art field map estimation software should produce the same reliable distortion correction observed with *Synth*. Two factors may potentially contribute to the variable performance of this approach. The tissue contrast properties of EPI data can vary significantly depending on image acquisition parameters. Therefore, one possibility is that SynB0-DisCo’s deep learning model may generalize poorly to unseen datasets whose resolution and contrast are quite different from its training data. A second possibility is that synthetic images produced by deep learning models are prone to introducing spurious artifacts into their outputs [22, 23]. The presence of such artifacts may be a significant source of registration variability. Both of these effects may have contributed to reducing SynB0-DisCo’s performance in the ABCD and MSC datasets. *Synth*’s ability to generate effective synthetic images based solely on the participant’s anatomical data without relying on a large training dataset avoids both of these potential issues by flexibly adapting to the contrast properties of a specific acquisition and minimizing the possibility of spurious artifacts.

While our results indicate that *Synth* is able to correct distortion comparably well to current field map based techniques, it is possible that advances in theory or software may improve field map-based corrections even further so that they reliably exceed *Synth*’s performance. We therefore do not advocate abandoning the collection of field map data entirely. Rather we propose that *Synth* distortion correction is a reliable substitute for field maps at the present time and given the current state of the art. We can unreservedly recommend using *Synth* as an effective tool for correcting distorted fMRI data when no field map data is available.

### 4.5. Future directions for field map-less distortion correction

Although the distortion corrections produced by *Synth* are high quality, several open questions remain relating to how this general approach can be implemented most effectively. Chief among these is whether *Synth*’s performance can be augmented further by including existing field map information derived using traditional approaches. The focus of the present report is assessing the quality of the field map corrections produced by *Synth* in a situation in which there is no initial estimate of the EPI image distortion provided to the underlying nonlinear warping software. Including an existing field map as an initial estimate may further increase *Synth*’s performance. Future iterations of *Synth* will include this ability and may produce even greater correction fidelity between functional and anatomical images than presented here.

*Synth* provides a high degree of flexibility, allowing the user to specify a custom linear model consisting of both main-effect terms and interaction terms between a user-specified number of source images, each decomposable into a user-specified number of RBF components. In general, more complex models will allow *Synth* to capture more complicated relationships between the voxel intensities in T1w/T2w images and the BOLD images. This flexibility comes at the cost of increased memory requirements and computational time, but can produce more accurate synthetic images that may ultimately improve registration quality. In our results, the RBF model parameters were chosen to produce a synthetic image of qualitatively sufficient visual similarity to their associated BOLD images. By default, *Synth* uses the RBF model parameters optimized for the ABCD and MSC datasets (i.e., a 12 component decomposition on each of the participants’ T1w/T2w images, along with pairwise interaction terms between T1w/T2w components). For users looking for guidance on selecting model specifications, our results indicate that synthetic images with contrast similarity (linear correlation coefficient) of with the target BOLD image are sufficient for producing high quality corrections. Additional exploration will be required to determine the optimal balance of time and quality.

Even though *Synth*’s underlying RBF model provides a great degree of flexibility in fitting the functions that map T1w/T2w voxel intensities to BOLD image voxel intensities, other data collection parameters and preprocessing steps may improve the quality of the synthetic images or reduce the needed complexity of the RBF model. For example, adjusting echo times during data acquisition may produce images that can be more accurately modeled with fewer RBF components while still retaining sufficient sensitivity to BOLD contrast. Alternatively or in addition, performing contrast enhancing preprocessing of the anatomical and BOLD images so that the resulting function relating their voxel intensities can be modeled more efficiently by the chosen RBF model may allow for simpler RBF models to perform equivalently. In our own preliminary tests and for the results reported here, we observed that synthetic images were more accurate when a bias field was estimated and removed from the anatomical images and image contrast was increased through histogram normalization (see Section 6.1.1 in Supplemental Methods).

### 4.6. Conclusion

The results reported here have demonstrated that it is possible to achieve high-quality EPI distortion correction without the need to collect separate field map data. We have shown that field map-less approaches, such as *Synth*, can perform comparably to existing gold-standard distortion correction approaches, and in some cases, may surpass them on measures of global and local image alignment quality. Removing the reliance on field maps to correct EPI distortion will allow researchers to recover samples with missing or corrupted field maps while maintaining high quality alignment between their anatomical and BOLD images. Field map-less distortion correction may prove to be a particular asset to researchers studying high motion cohorts, such as pediatric or neuropsychiatric populations [55], where acquiring high quality field map data is a significant challenge. Reliable field map-less distortion correction shows great promise for overcoming the limitations arising from the acquisition, processing, and quality checking of field map data and has the potential to greatly simplify data processing in the neuroimaging field.

## Code sharing

The *Synth* software package and other utilities and scripts used for this project can be downloaded at https://gitlab.com/vanandrew/omni.

## Data availability

The ABCD data repository grows and changes over time. The ABCD data used in this report came from ABCD collection 3165 and the Annual Release 2.0, DOI 10.15154/1503209.

The MSC dataset used in this report can downloaded at can be found on OpenNeuro.org, DOI 10.18112/open-neuro.ds000224.v1.0.3.

## Acknowledgements

This work was supported by DA007261 (D.F.M.), MH096773 (D.A.F, N.U.F.D.), MH122066 (D.A.F., N.U.F.D.), MH121276 (D.A.F., N.U.F.D), MH124567 (D.A.F., N.U.F.D.), MH100019 (T.O.L.), NS110332 (D.J.N., N.U.F.D.), NS088590 (N.U.F.D.), Kiwanis Neuroscience Research Foundation (N.U.F.D.), the Jacobs Foundation grant 2016121703 (N.U.F.D.).

## ABCD acknowledgement

Data used in the preparation of this article were obtained from the Adolescent Brain Cognitive Development (ABCD) Study (https://abcdstudy.org), held in the NIMH Data Archive (NDA). This is a multisite, longitudinal study designed to recruit more than 10,000 children age 9-10 and follow them over 10 years into early adulthood. The ABCD Study is supported by the National Institutes of Health and additional federal partners under award numbers U01DA041022, U01DA041028, U01DA041048, U01DA041089, U01DA041106, U01DA041117, U01DA041120, U01DA041134, U01DA041148, U01DA041156, U01DA041174, U24DA041123, U24DA041147, U01DA041093, and U01DA041025. A full list of supporters is available at https://abcdstudy.org/federal-partners.html. A listing of participating sites and a complete listing of the study investigators can be found at https://abcdstudy.org/scientists/workgroups/. ABCD consortium investigators designed and implemented the study and/or provided data but did not necessarily participate in analysis or writing of this report. This manuscript reflects the views of the authors and may not reflect the opinions or views of the NIH or ABCD consortium investigators.

## 5. Supplemental Material

### 5.1. Scanner manufacturers and models comprising the ABCD dataset

### 5.2. Supplemental Figures

**Supplemental Figure 1:**
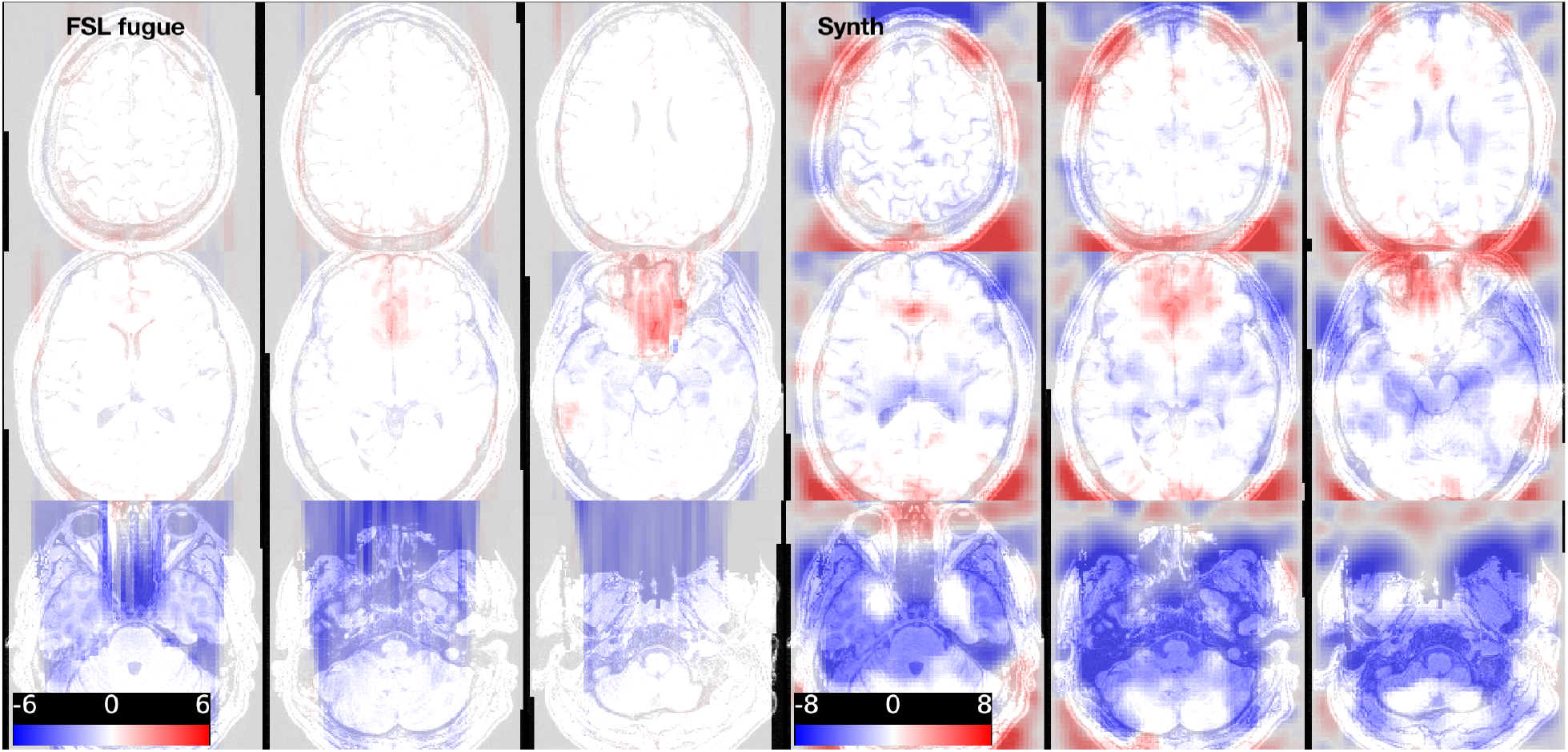
Comparison of *Synth* and FSL *fugue* field map correction for randomly selected MSC participant. **(Left)** 3×3 panel depicts the displacement field estimated by FSL *fugue* for a single MSC participant **(Right)** 3×3 panel depicts the displacement field estimated for the same participant by *Synth*. FSL *fugue* estimates the distortion correcting warp within the bounds of a tightly cropped brain mask, yielding warps that only affect a relatively compact region of the image that contains the brain. *Synth* estimates the warps over the entire field of view and therefore produces field map corrections in which more voxels experience shifts. Overall displacement magnitudes are smaller in this session for FSL *fugue* warps. Depicted voxel shifts are in millimeter units.

**Supplemental Figure 2:**
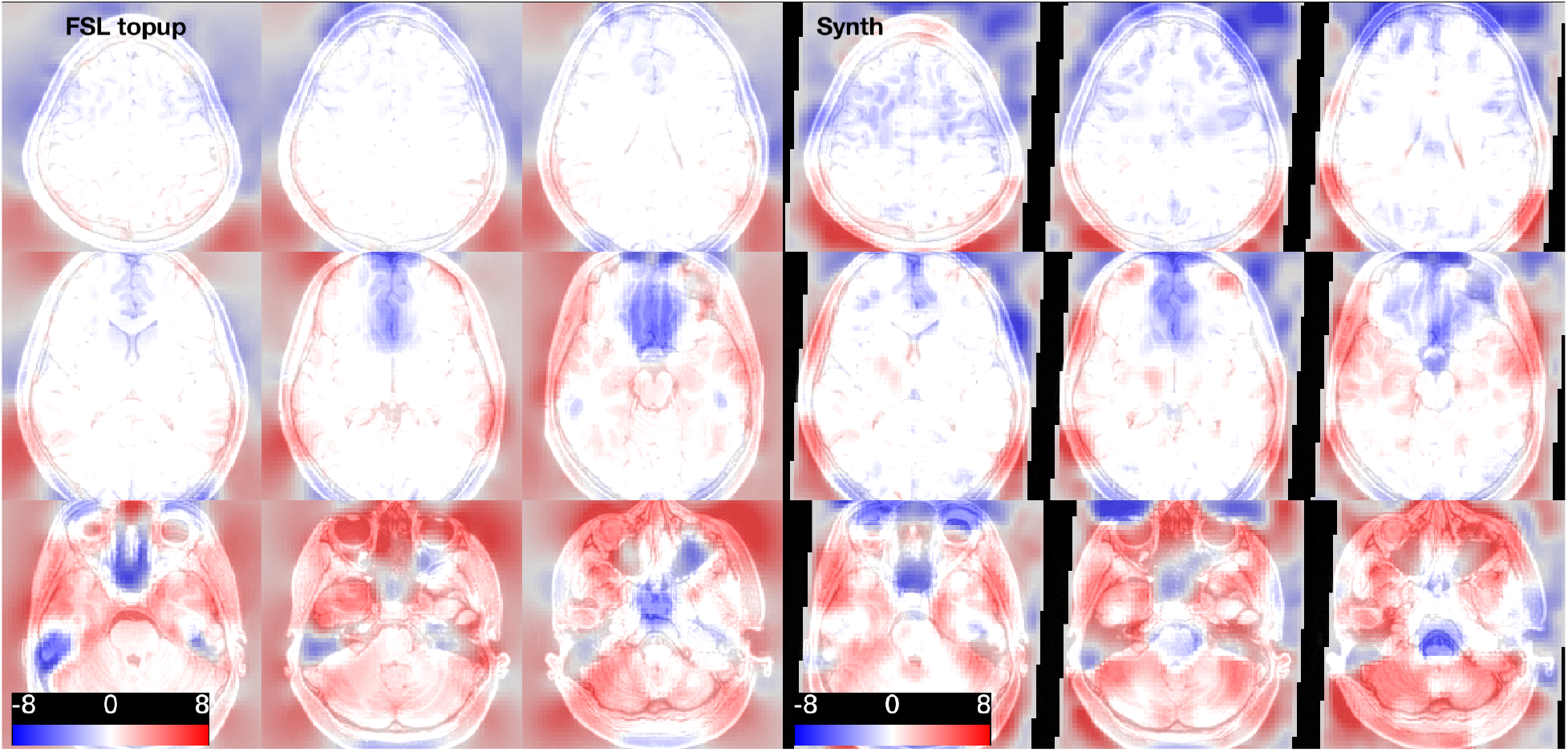
Comparison of *Synth* and FSL *topup* field map correction for randomly selected ABCD participant. **(Left)** 3×3 panel depicts the displacement field estimated by FSL *topup* for a single ABCD participant **(Right)** 3×3 panel depicts the displacement field estimated for the same participant by *Synth*. Like *Synth*, FSL *topup* estimates the distortion correcting warp over the entire field of view. Both *Synth* and FSL *topup* produce warps that are qualitatively similar to one another. Depicted voxel shifts are in millimeter units.

**Supplemental Figure 3:**
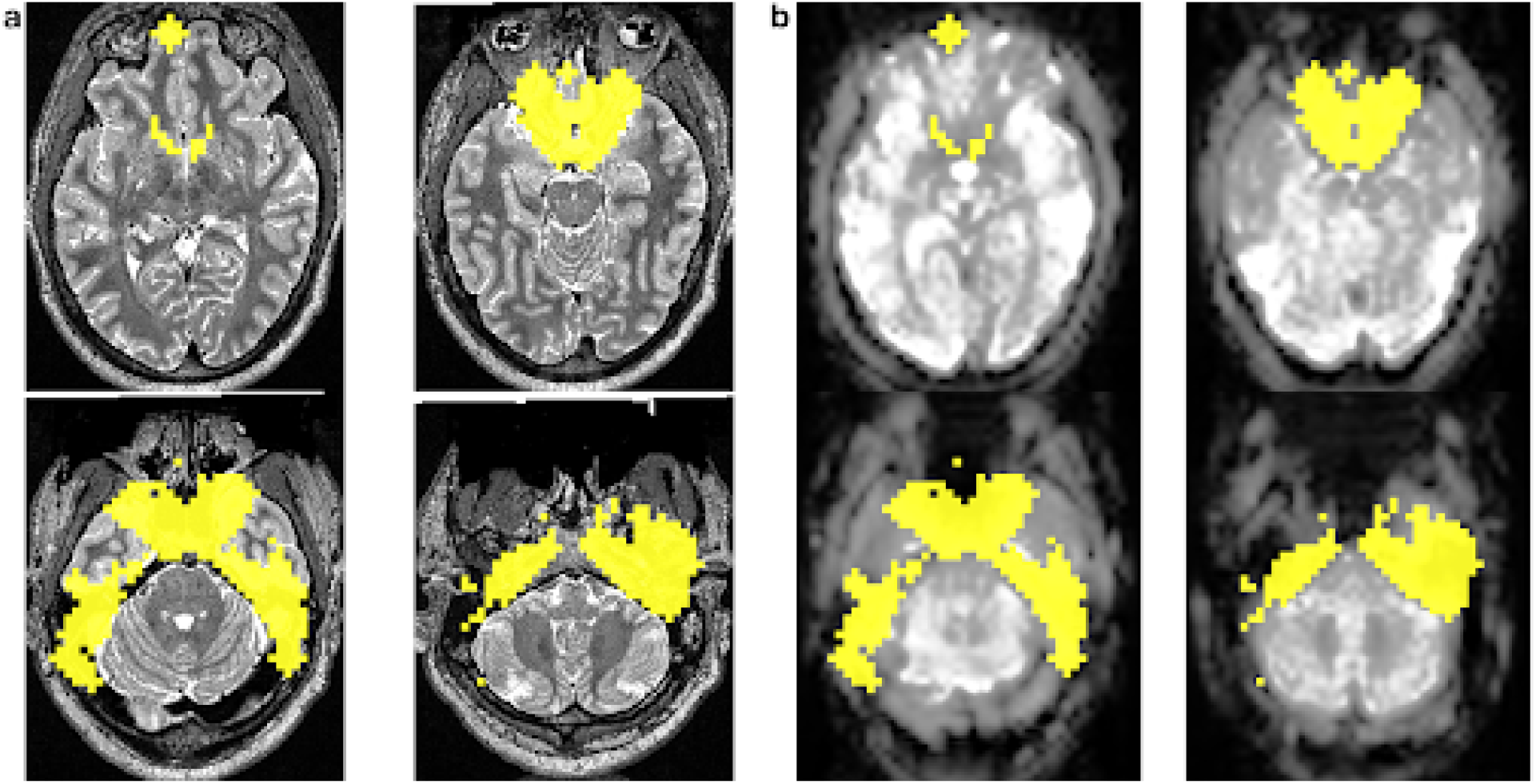
Representative signal-to-noise exclusion mask. **(a)** Example participant T2w images (underlay) depicting slices that typically contain regions of significant signal loss in EPI scans. **(b)** The corresponding closely aligned slices from the participant’s EPI data. In both sets of images, the yellow overlay depicts regions of extreme signal loss detected with the LDA procedure outlined above. In these regions of extreme signal loss, no correspondence exists between a participants’ anatomical and BOLD data. Therefore, constructing the synthetic BOLD image, the contributions of voxels in this region are down weighed when running *Synth*. Note qualitative similarity between the noise mask detected with this method and group average map of fMRI signal loss reported in [56].

### 5.3. Distortion Correction with Synth

#### Algorithm 1

Distortion Correction with *Synth*

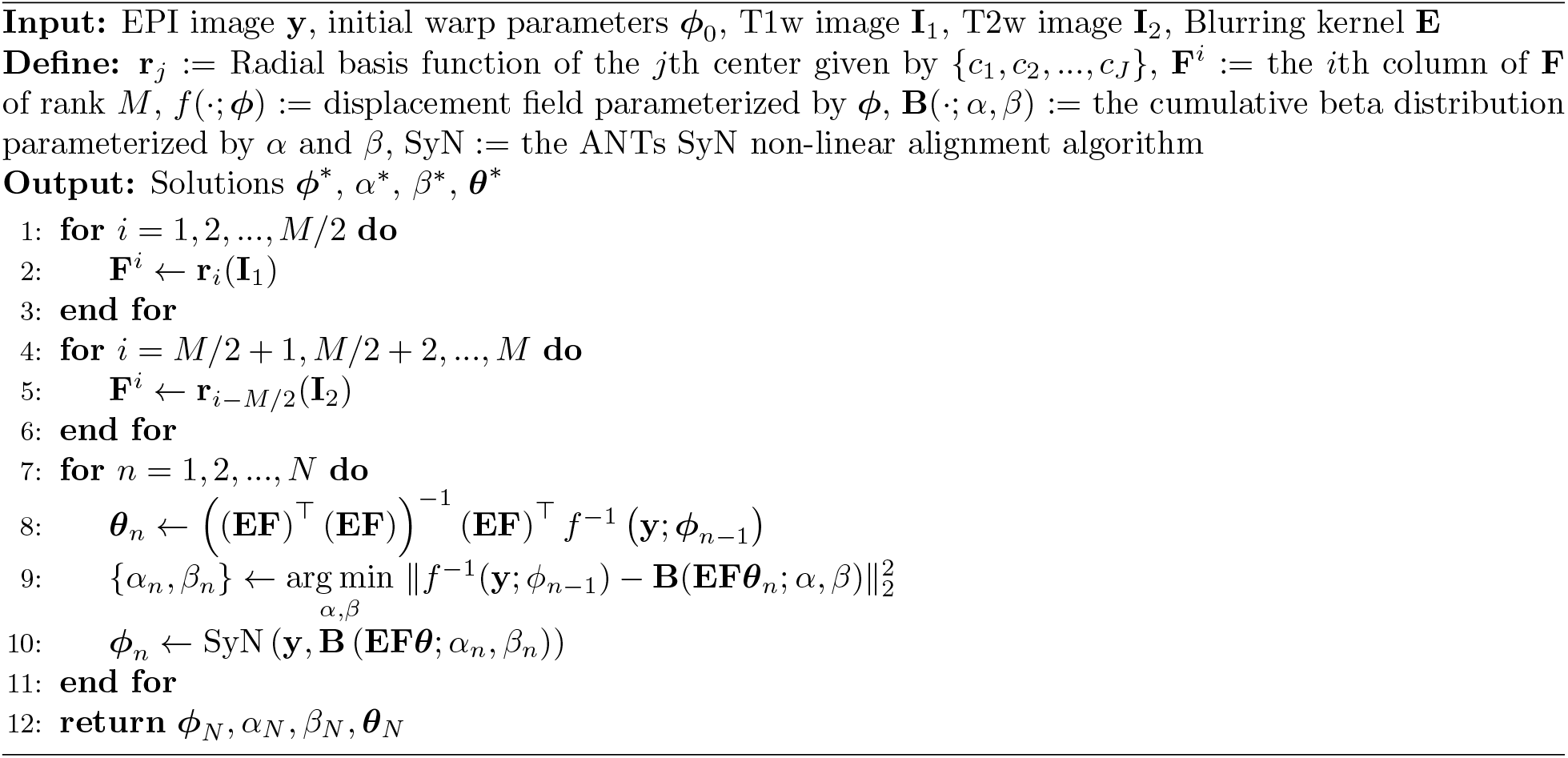

### 5.4. Statistical tables for global and local similarity metrics (ABCD dataset)

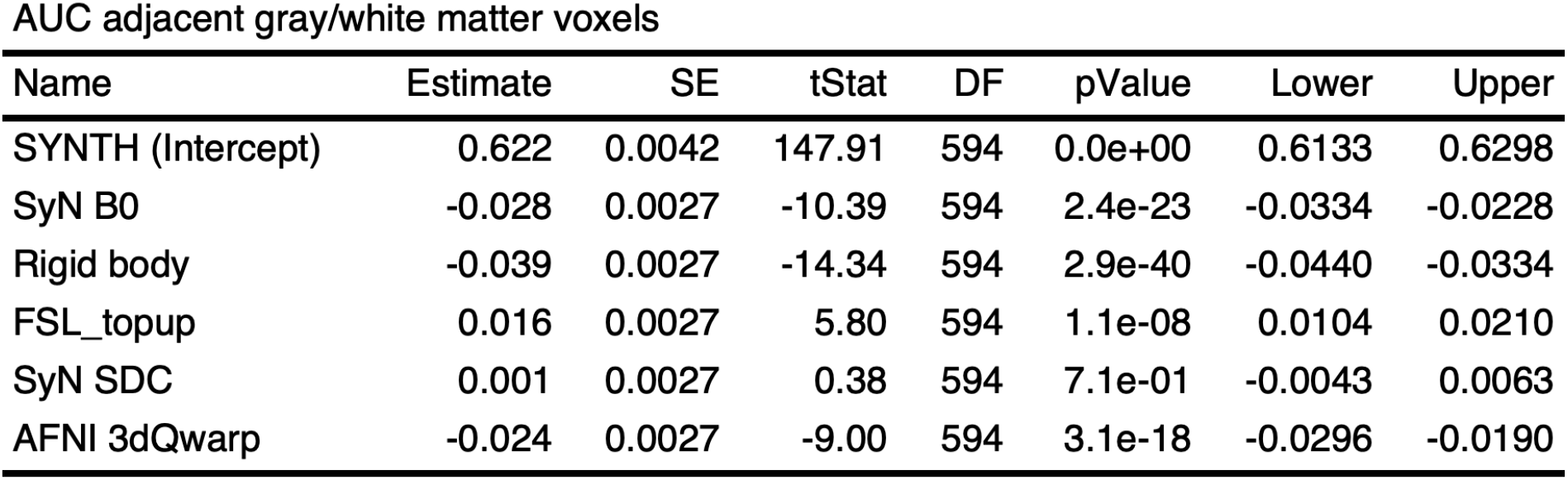

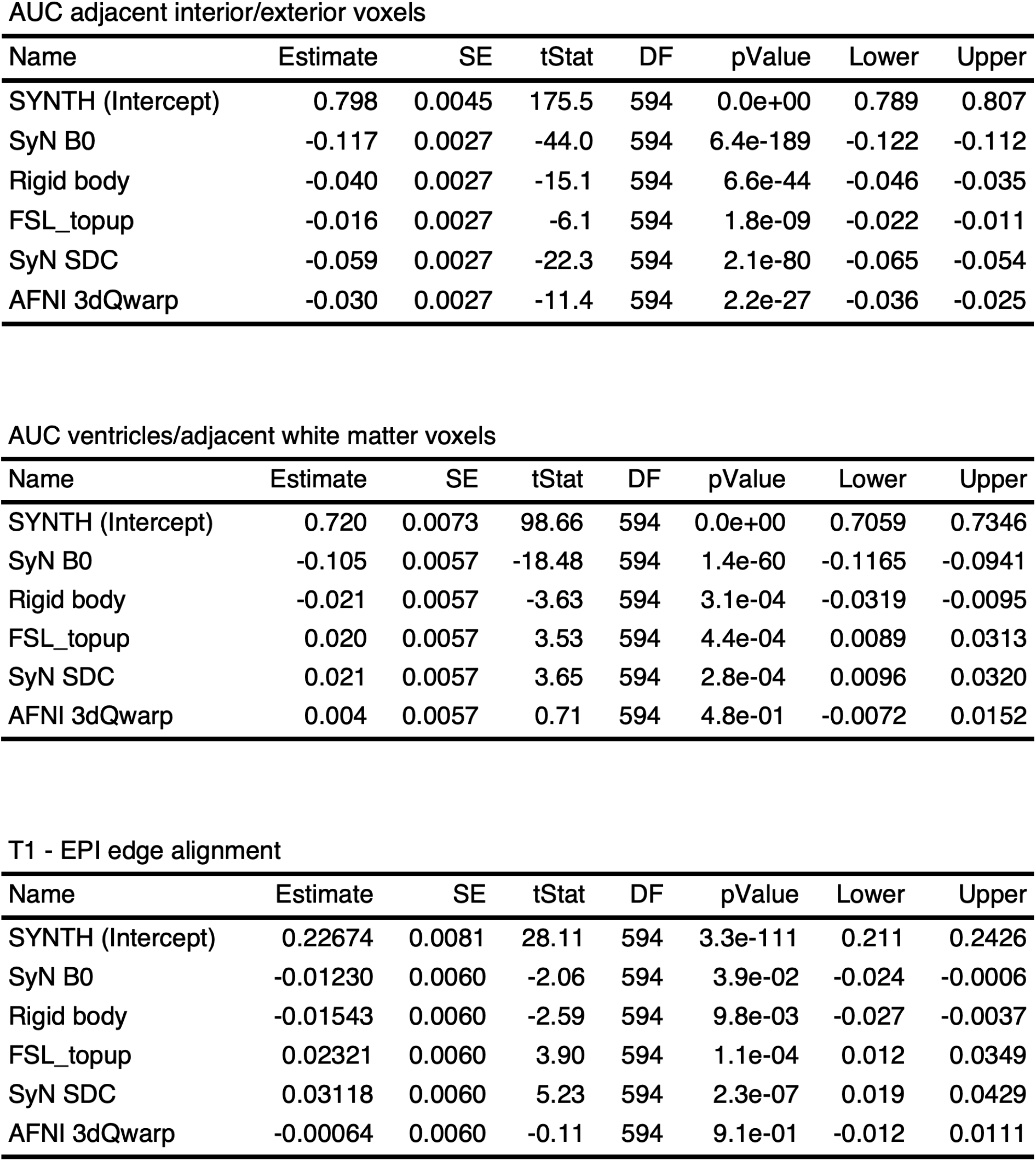

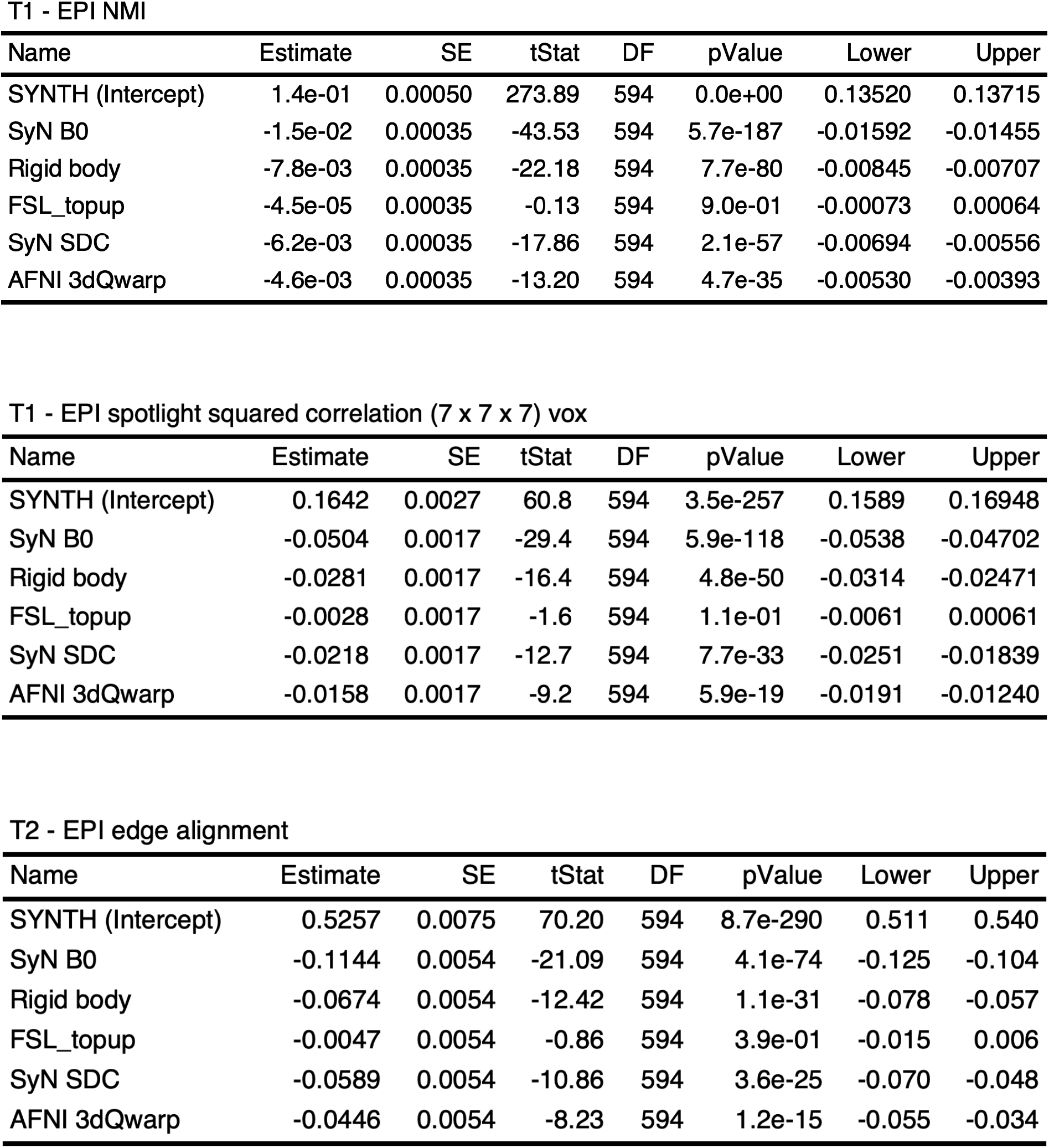

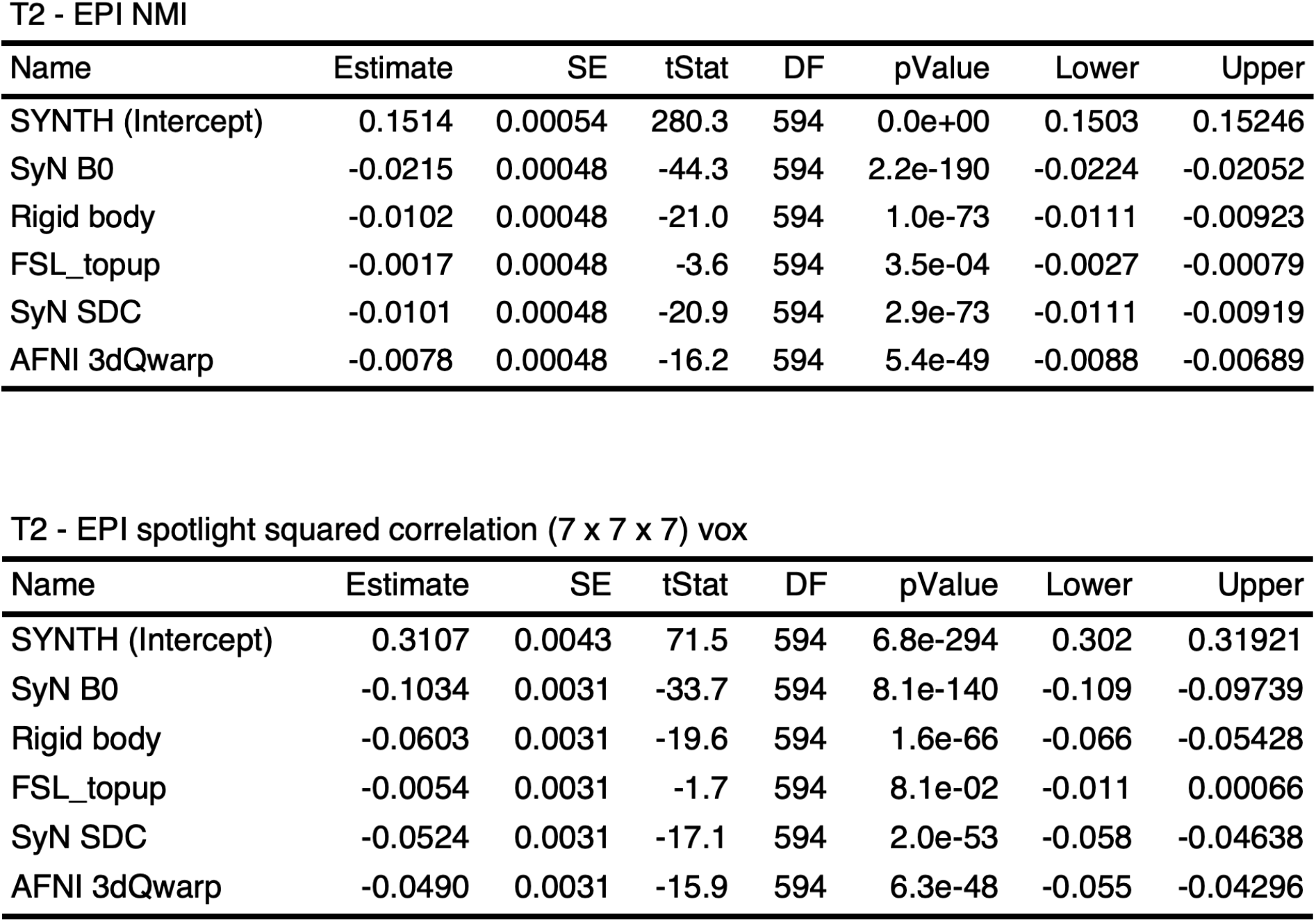

### 5.5. Statistical tables for global and local similarity metrics (MSC dataset)

5.5. Statistical tables for global and local similarity metrics (MSC dataset) AUC adjacent gray/white matter voxels
| Name | Estimate | SE | tStat | DF | pValue | Lower | Upper |
| --- | --- | --- | --- | --- | --- | --- | --- |
| SYNTH (Intercept) | 0.5945 | 0.0082 | 72.6 | 495 | 4.9e-266 | 0.5784 | 0.611 |
| SyN B0 | -0.0533 | 0.0023 | -23.4 | 495 | 3.5e-82 | -0.0578 | -0.049 |
| Rigid body | -0.0217 | 0.0023 | -9.5 | 495 | 7.0e-20 | -0.0262 | -0.017 |
| FSL_topup | 0.0079 | 0.0023 | 3.5 | 495 | 5.8e-04 | 0.0034 | 0.012 |
| SyN SDC | -0.0392 | 0.0023 | -17.2 | 495 | 1.6e-52 | -0.0437 | -0.035 |

AUC adjacent interior/exterior voxels
| Name | Estimate | SE | tStat | DF | pValue | Lower | Upper |
| --- | --- | --- | --- | --- | --- | --- | --- |
| SYNTH (Intercept) | 0.723 | 0.0103 | 70.4 | 495 | 4.5e-260 | 0.703 | 0.744 |
| SyN B0 | -0.045 | 0.0022 | -20.0 | 495 | 1.3e-65 | -0.049 | -0.040 |
| Rigid body | -0.019 | 0.0022 | -8.4 | 495 | 4.4e-16 | -0.023 | -0.014 |
| FSL_topup | -0.022 | 0.0022 | -9.9 | 495 | 4.6e-21 | -0.027 | -0.018 |
| SyN SDC | -0.058 | 0.0022 | -25.8 | 495 | 9.8e-94 | -0.062 | -0.053 |

AUC ventricles/adjacent white matter voxels
| Name | Estimate | SE | tStat | DF | pValue | Lower | Upper |
| --- | --- | --- | --- | --- | --- | --- | --- |
| SYNTH (Intercept) | 0.7634 | 0.0170 | 44.81 | 495 | 2.6e-176 | 0.730 | 0.7969 |
| SyN B0 | -0.0260 | 0.0045 | -5.72 | 495 | 1.8e-08 | -0.035 | -0.0171 |
| Rigid body | -0.0611 | 0.0045 | -13.43 | 495 | 2.7e-35 | -0.070 | -0.0522 |
| FSL_topup | -0.0044 | 0.0045 | -0.96 | 495 | 3.4e-01 | -0.013 | 0.0046 |
| SyN SDC | -0.0820 | 0.0045 | -18.03 | 495 | 3.0e-56 | -0.091 | -0.0731 |

T1 - EPI edge alignment
| Name | Estimate | SE | tStat | DF | pValue | Lower | Upper |
| --- | --- | --- | --- | --- | --- | --- | --- |
| SYNTH (Intercept) | 0.395 | 0.0111 | 35.5 | 495 | 3.4e-138 | 0.373 | 0.417 |
| SyN B0 | -0.073 | 0.0047 | -15.4 | 495 | 5.9e-44 | -0.082 | -0.064 |
| Rigid body | -0.034 | 0.0047 | -7.2 | 495 | 2.8e-12 | -0.043 | -0.025 |
| FSL_topup | -0.023 | 0.0047 | -4.8 | 495 | 2.0e-06 | -0.032 | -0.013 |
| SyN SDC | -0.117 | 0.0047 | -24.7 | 495 | 3.6e-88 | -0.126 | -0.108 |

T1 - EPI NMI
| Name | Estimate | SE | tStat | DF | pValue | Lower | Upper |
| --- | --- | --- | --- | --- | --- | --- | --- |
| SYNTH (Intercept) | 0.1434 | 0.00134 | 107 | 495 | 0.0e+00 | 0.1408 | 0.1460 |
| SyN B0 | -0.0143 | 0.00036 | -40 | 495 | 4.3e-158 | -0.0150 | -0.0136 |
| Rigid body | -0.0102 | 0.00036 | -29 | 495 | 3.8e-107 | -0.0109 | -0.0095 |
| FSL_topup | -0.0044 | 0.00036 | -12 | 495 | 4.9e-31 | -0.0051 | -0.0037 |
| SyN SDC | -0.0159 | 0.00036 | -45 | 495 | 1.5e-175 | -0.0166 | -0.0152 |

T1 - EPI spotlight squared correlation (7 x 7 x 7) vox
| Name | Estimate | SE | tStat | DF | pValue | Lower | Upper |
| --- | --- | --- | --- | --- | --- | --- | --- |
| SYNTH (Intercept) | 0.254 | 0.0061 | 41.7 | 495 | 3.6e-164 | 0.242 | 0.266 |
| SyN B0 | -0.068 | 0.0020 | -33.8 | 495 | 1.6e-130 | -0.072 | -0.064 |
| Rigid body | -0.040 | 0.0020 | -20.1 | 495 | 4.9e-66 | -0.044 | -0.036 |
| FSL_topup | -0.018 | 0.0020 | -9.2 | 495 | 1.2e-18 | -0.022 | -0.015 |
| SyN SDC | -0.069 | 0.0020 | -34.2 | 495 | 2.4e-132 | -0.073 | -0.065 |

T2 - EPI edge alignment
| Name | Estimate | SE | tStat | DF | pValue | Lower | Upper |
| --- | --- | --- | --- | --- | --- | --- | --- |
| SYNTH (Intercept) | 0.48896 | 0.0092 | 53.25 | 495 | 4.9e-207 | 0.4709 | 0.507 |
| SyN B0 | -0.07349 | 0.0053 | -13.93 | 495 | 1.8e-37 | -0.0839 | -0.063 |
| Rigid body | -0.04319 | 0.0053 | -8.19 | 495 | 2.2e-15 | -0.0536 | -0.033 |
| FSL_topup | 0.00078 | 0.0053 | 0.15 | 495 | 8.8e-01 | -0.0096 | 0.011 |
| SyN SDC | -0.13181 | 0.0053 | -24.99 | 495 | 9.1e-90 | -0.1422 | -0.121 |

T2 - EPI NMI
| Name | Estimate | SE | tStat | DF | pValue | Lower | Upper |
| --- | --- | --- | --- | --- | --- | --- | --- |
| SYNTH (Intercept) | 0.1478 | 0.00107 | 138 | 495 | 0.0e+00 | 0.1457 | 0.1499 |
| SyN B0 | -0.0136 | 0.00035 | -39 | 495 | 2.6e-154 | -0.0142 | -0.0129 |
| Rigid body | -0.0096 | 0.00035 | -28 | 495 | 4.0e-103 | -0.0103 | -0.0089 |
| FSL_topup | -0.0035 | 0.00035 | -10 | 495 | 5.9e-22 | -0.0042 | -0.0028 |
| SyN SDC | -0.0164 | 0.00035 | -48 | 495 | 1.3e-186 | -0.0171 | -0.0157 |

T2 - EPI spotlight squared correlation (7 x 7 x 7) vox
| Name | Estimate | SE | tStat | DF | pValue | Lower | Upper |
| --- | --- | --- | --- | --- | --- | --- | --- |
| SYNTH (Intercept) | 0.24098 | 0.0055 | 44.20 | 495 | 5.8e-174 | 0.2303 | 0.2517 |
| SyN B0 | -0.04729 | 0.0022 | -21.07 | 495 | 7.8e-71 | -0.0517 | -0.0429 |
| Rigid body | -0.03714 | 0.0022 | -16.55 | 495 | 2.9e-49 | -0.0415 | -0.0327 |
| FSL_topup | -0.00095 | 0.0022 | -0.42 | 495 | 6.7e-01 | -0.0054 | 0.0035 |
| SyN SDC | -0.06795 | 0.0022 | -30.27 | 495 | 1.0e-114 | -0.0724 | -0.0635 |

## 6. Supplemental Methods

### 6.1. Description of common pipeline

Distortion correction methods were compared using a common registration pipeline, which differed only in the distortion correction method and functional-to-T1w alignment. This includes alignment of anatomical images to atlas, functional framewise alignment and resampling of the final aligned functional data to 3 mm. Details of each common step are explained in the following sections.

#### 6.1.1. Anatomical registration/segmentation

Prior to registration, both the T1w and T2w images were bias field corrected using *N4BiasFieldCorrection* [57] (spline distance = 100, initial mesh resolution = 1×1×1). Anatomical alignment was accomplished by extracting edge images from each participants’ T1w and T2w images using AFNI’s 3dedge utility and aligned with a rigid body transform using FLIRT and a correlation ratio cost function. The resulting transformation parameters were saved for later use.

To improve the reliability of the skullstripping procedure, an intermediate anatomical image was utilized. This image was created using the following procedure: 1) T1w and T2w images were rescaled so that values were between 0 and 1; 2) the intermediate volume was calculated where the value at each voxel is determined by the square root of the sum of squares of the corresponding voxels in the scaled T1w and T2w images. The resulting image was passed into BET [58] (fractional intensity threshold = 0.1) for skullstripping. The skullstripping mask was then applied to the participant’s aligned T1w and T2w images.

For each participant, we aligned their debiased, skullstripped T1w volume to a common template, the TRIO Y NDC atlas, using a 9-parameter affine transformation estimated with FLIRT and a mutual information cost function. We then combined the resulting transformations to align both T1w and T2w volumes to the TRIO atlas using windowed sinc-function interpolation. Anatomical segmentation was done using recon-all from FreeSurfer 6.0 (Fischl, 2012) with the -T2 flag enabled.

#### 6.1.2. Alignment of EPI images

Framewise alignment of EPI images was accomplished using AFNI’s *3dAllineate*. First, a reference image was constructed by averaging 100 aligned frames from the resting state times series taken from time intervals with the lowest DVARs values [59]. To determine these intervals, a DVARs time series was calculated by computing the temporal derivative of each voxel’s time series and normalizing its standard deviation to be equal to 1. For each frame, a DVARs value was calculated as the spatial mean of the absolute value of the derivative time series. Because we were interested in extracting the frames to construct the reference EPI image from prolonged intervals of relatively low DVARs values, we then low-pass filtered the DVARs time series by bi-directionally filtering it with a 1st order digital Butterworth filter with normalized cut-off frequency of 0.16. To create the reference image, we selected 100 frames associated with the lowest smoothed DVARs values and aligned them to the individual frame that had the lowest DVARs value in the session using rigid body transformations. Alignment parameters were estimated with *3dAllineate* and employed a least-squares cost function along with the -autoweight option. Finally we constructed a reference image for the session by averaging the 100 aligned frames together.

Lastly, framewise alignment parameters were estimated by registering each frame individually to the reference image. To reduce the influence of signal drift on framewise registration and to increase the contribution of properly aligned edges to the cost function, we spatially high-pass filtered the reference functional image and each functional frame independently by smoothing them with large 12 mm FWHM gaussian kernel and subtracting result from the original image. These high-pass filtered volumes were then rigid body aligned using *3dAllineate* which optimized the cost function calculated over 75% of the voxels.

#### 6.1.3. Resting state denoising

Resting state data was denoised by regressing a set of nuisance covariates from each voxel’s time series. The design matrix included a constant term; a linear trend component; the first 8 low order Legendre polynomials -to remove low frequency fluctuations; time series corresponding to the 6 framewise rigid-body alignment parameters; and the first derivative of each of the motion parameters along with two additional lagged copies. In addition, the design matrix included nuisance white matter time series as well as time series from nearby voxels outside of the brain used to model global and regional nuisance variability [60, 61, 62]. The white matter regressors were constructed by extracting and normalizing time series from all voxels labeled as white matter by the participant’s FreeSurfer parcellation. Singular value decomposition was then applied to extract 12 time series accounting for the most white matter variance. A similar procedure was used to construct the external signal nuisance regressors. Here, time series from voxels residing within a 5 voxel shell surrounding the brain were extracted. Again, the first 12 singular vectors were computed and included in the denoising design matrix. The complete set of nuisance variables was then projected from the voxel time series using linear regression. Lastly, the regression model residuals were bandpass filtered to retain only frequencies between 0.001 and 0.1 Hz. Frame censoring to remove high motion frames was employed using a weighted correlation approach [63, 59, 64]. First, we computed a DVARs time series across all brain matter voxels. Frames in which DVARs exceeded two standard deviations were flagged and given a zero weighting. To reduce the influence of any contamination remaining in adjacent frames, the weighting of the three frames preceding and following were also reduced in proportion to their absolute distance from the censored frame.

### 6.2. Description of radial basis function model used to estimate the synthetic EPI image

*Synth* allows the user to specify a regression model that maps source image intensities to target image intensities. The radial basis components were constructed using the following procedure: Let **I**_*m*_ refer to an individual source image, e.g.,T1w or T2w; and {*c*_1_, *c*_2_, …, *c*_*J*_ ∈ *C*} refers to a set of *J* evenly spaced values that lie between the maximum and minimum values of **I**_*m*_. Then each radial basis component, **r**_*j*_, of an image, **I**_*m*_, is constructed according to the following equation:

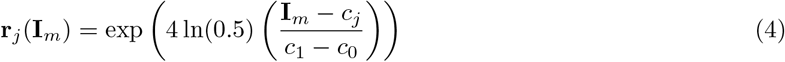

A visual representation of the images produced by applying Eqn. 4 to T1w and T2w images is depicted in Figure 1a. The final design matrix fit by *Synth* consists of each radial basis component concatenated into a single matrix **F**, along with intercept terms, **e**_*m*_, (a vector of all ones) for each source image, such that:

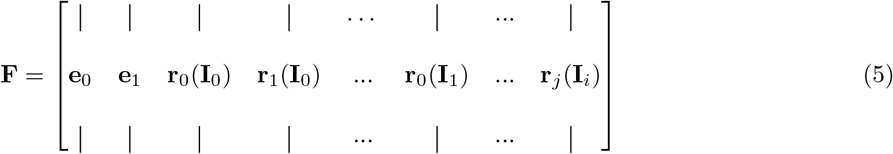

For both datasets, **F** was constructed from a 12 component decomposition on each of the participants’ T1w/T2w images, along with all T1w/T2w pairwise interaction terms. As in traditional linear models, these interaction terms are simply the pairwise product of the columns of **F**.

### 6.3. The blur operator

While *Synth* estimates the field map corrections at the 1 mm isotropic voxel resolution of the T1w/T2w and synthetic EPI images, the true EPI images are acquired at much lower resolutions. In the case of the MSC dataset, this corresponds to a 4 mm native voxel resolution resampled to 1 mm isotropic resolution; and for the ABCD dataset this corresponds to 2.6 mm native resolution resampled to 1 mm. When lower resolution EPI images are upsampled to the higher resolution grid, their lower resolution manifests as apparent blur. *Synth* models this blur (**E** in Eqn. 1) as the convolution of a source image with an Epanachnikov kernel, whose bandwidth is related to the resolution difference between the EPI and anatomical images. This kernel is given by the following expression:

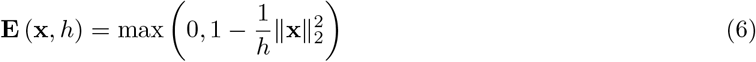

where **x** ∈ ℝ^3^ is a location on the kernel, and *h* is the bandwidth parameter. For the results in this paper, the bandwidths of the Epanechnikov kernels were set to *h* = 6 and *h* = 10 for the ABCD and MSC dataset respectively.

*Synth* models the difference in effective spatial resolution between T1w/T2w and BOLD images by blurring the radial basis function (RBF) components of the anatomical images with an Epanachnikov smoothing kernel prior to fitting the model to the target BOLD image. In principle, the optical blurring should occur only after the field map deformation is applied, rather than prior to it. However, implementing the model in this way would require repeatedly blurring the high spatial resolution radial basis components of the T1w/T2w source images after every iterative update of the estimated field map parameters. A naive implementation of this approach would be computationally prohibitive. Prior research has shown that in practice the sort of approximation implemented in *Synth* in many cases does not introduce a significant amount of error [65]. However, while the quality of the field map corrections produced by *Synth* appears to be quite high, some improvement may yet be gleaned by using the “correct model”, assuming the technical challenges to its implementation can be surmounted.

### 6.4. Contrast correction for error induced image bias

When source and target images are acquired at substantially different spatial resolutions, or in the early phases of alignment when source and target images are far out of register, the synthetic images produced by *Synth* exhibit significantly “flattened” contrast, even if the rank order voxel intensities comprising the synthetic image largely correspond to those observed in the target image. This is due to the fact that errors in registration or large differences between source and target image resolution bias estimates of the radial basis function parameters, ***θ***, toward the target image mean. In an extreme, but illustrative, case where image registration is so far off that there is no meaningful relationship between corresponding source and target image voxels, the synthetic image will simply be a volume in which all voxels have the same intensity as the target image mean. That is, it will have perfectly ‘flat’ contrast. As registration improves some, the synthetic image will begin to exhibit more accurately the contrast properties of the target image. In general, however, the contrast of naively estimated synthetic images will tend to be somewhat flattened due to error. To address this, *Synth* includes a tone curve adjustment option that increases contrast of the final synthetic image. This is accomplished by passing the synthetic image voxel intensities through a monotonic non-linear curve, a process similar to tone curve adjustment in traditional image processing. The curve is constrained to reside within the set of beta function cumulative distributions -distributions well suited for modeling tone curves that tend to be sigmoidal- and its parameters, *α* and *β*, are optimized by minimizing the negative linear correlation coefficient between the target and synthetic images.

### 6.5. Estimating field maps with an undistorted synthetic EPI image

The warp parameters that map the undistorted synthetic EPI image to the reference EPI image were estimated using a three step iterative procedure. The final iteration of this procedure produced the estimated field map corrections used for our analyses. This procedure was conducted as follows:

1. Using *Synth*, construct an initial synthetic EPI image based on the current best affine alignment between T1w, T2w and a reference EPI volume. An EPI signal-to-noise mask was used to reduce the contribution of low signal-to-noise areas when constructing the synthetic EPI (see Section 6.6).
2. Estimate a warp that maps the current synthetic EPI to the reference EPI image. With the exception of the first iteration, each warp estimate is initialized from the warp estimate of the previous iteration.
3. Apply the inverse of this warp, which approximates a field map correction, to the reference EPI image.
4. Repeat steps 1-3, at each iteration, using *Synth* to re-estimate the synthetic EPI image using the most recently corrected EPI image (see Eqn. 1).

For the results presented here, we estimated the field maps using the ANTs SyN algorithm. Each subsequent iteration used a smaller gradient step size, 3, 1, 0.1 for each iteration respectively. SyN parameters were set to use a neighborhood cross-correlation cost function (radius = 2), update and total field variance parameters of 0, winsorize limits of [0.005, 0.995], histogram matching, BSpline interpolation of order 5, convergence parameters of 500×0, 1e-6 and 15, smoothing parameters of 0×0, and shrink factor parameters 2×1. In order to leverage the detailed spatial information available in the high resolution synthetic EPI images during the estimation of the field map, the warps at each iteration were constructed in the 1mm × 1mm × 1mm native resolution of the T1w/T2w source images.

### 6.6. Creating the EPI signal-to-noise mask

To construct a binary mask that would serve to label EPI voxels suffering from a high degree of noise or signal dropout we implemented a linear discriminant analysis procedure. We began by using the brain mask derived from the skullstripping stage of the anatomical volume and aligned it to the functional dataset using the affine transformation generated from *Synth*. This mask served as an initial guess, or prior estimate, for which voxels were likely to represent meaningful fMRI signals in the resting state EPI dataset, and those which represent noise and areas of signal dropout Supplemental Figure 3. Using this initial labeling, we implemented a between-class linear discriminant analysis (LDA) based, first, on the raw un-centered and un-scaled voxel time series and then computed the projection of each voxel’s time series onto the resulting principal discriminant vector. We repeated this LDA procedure on the centered and normalized time series from each voxel, again projecting the times series back onto the principal discriminant vector. Thus, each voxel was associated with two projection values. We then created a third value associated with each voxel by computing the product of the two principal discriminant vector projections. A second level of LDA was done on these three values. Lastly, Otsu’s method [66] was used to label each voxel as ‘signal’ or ‘noise’, depending upon its projection onto this final 3-element principal discriminant vector. We refined this labeling by repeating the entire LDA procedure over 20 iterations. During each subsequent iteration the labels produced by the previous iteration were used during the LDA procedure. Sufficiently iterated, this approach converges to a stable binary signal-or-noise label for each voxel. Lastly, to construct a mask containing only within-brain noise voxels —voxels most likely to be affected by signal dropout— we masked the final result of this iterative process to remove non-brain voxels (defined by the skullstrip mask) and morphologically open and dilate the image by 1 voxel. This mask was used as a weighting volume to further refine the estimate of the synthetic functional EPI by reducing the contributions of brain regions where no correspondence could exist between BOLD images and T1w/T2w images due to signal drop out.

### 6.7. Distortion correction using alternating descent optimization with Synth

Distortion correction with *Synth* is accomplished through an alternating descent optimization approach, which solves for each parameter individually while keeping the others fixed. At the end of each iteration of alternating descent, updates to the distortion correction estimation improves the accuracy of the synthetic image constructed during the subsequent iteration. These sequential improvements allow for iteratively more accurate distortion correction. The complete procedure is described in Algorithm 1.

Prior to the alternating descent procedure, initial warp parameters, ***ϕ***_0_, are initialized by a rigid-body transform that aligns the T2w anatomical image and BOLD functional image using a mutual information cost function. The rigid-body parameters from this step are then converted into a displacement field and applied to the BOLD image, aligning it to the anatomical images that underlie the matrix, **F**. Algorithm 1 begins by constructing the radial basis function matrix, **F**, from the T1w/T2w anatomical images. Then the main alternating minimization loop is run for each iteration *n*. First, ***θ***_*n*_ is computed by solving the linear system, *f* ^−1^ (**y**; ***ϕ***_*n*−1_) = **EF*θ***_*n*_. Here, *f* ^−1^ denotes the inverse displacement field that non-linearly transforms the real EPI image to the space of the synthetic EPI image. ***θ***_*n*_ is then used to construct an intermediate synthetic image. Next, global contrast parameters, *α*_*n*_ and *β*_*n*_ are estimated to improve overall similarity between the BOLD image and the intermediate synthetic image. Finally, a new estimate for the displacement field parameters, ***ϕ***_*n*_, are estimated using the ANTs SyN non-linear warping algorithm. Each iteration updates ***ϕ***_*n*_, ***θ***_*n*_, *α*_*n*_, and *β*_*n*_ until a specified number of iterations, *N*, is reached.

